# Unraveling wheat endosperm development: epigenetic regulation and novel regulators for enhanced yield and quality

**DOI:** 10.1101/2024.01.08.574643

**Authors:** Long Zhao, Jinchao Chen, Zhaoheng Zhang, Wenying Wu, Xuelei Lin, Mingxiang Gao, Yiman Yang, Peng Zhao, Yingyin Yao, Aiming Zhang, Dongcheng Liu, Dongzhi Wang, Jun Xiao

**Affiliations:** Key Laboratory of Plant Cell and Chromosome Engineering, Institute of Genetics and Developmental Biology, Chinese Academy of Sciences, Beijing 100101, China; University of Chinese Academy of Sciences, Beijing 100049, China; Hebei Agricultural University, Baoding, Hebei 071001, China; Nanjing Agricultural University, Nanjing, Jiangsu 210095, China; State Key Laboratory for Crop Stress Resistance and High-Efficiency Production, College of Agronomy, Northwest A&F University, Yangling 712100, China; China Agricultural University, Beijing 100193, China; Centre of Excellence for Plant and Microbial Science (CEPAMS), JIC-CAS, Beijing 100101, China

**Keywords:** wheat endosperm, starch, seed storage protein, TRN, epigenetic

## Abstract

Starch content and seed storage protein (SSP) composition are critical factors influencing wheat grain yield and quality. To uncover the molecular mechanisms governing their biosynthesis, we conducted transcriptome and epigenome profiling across key endosperm developmental stages, revealing that chromatin accessibility, H3K27ac, and H3K27me3 collectively regulate SSP and starch genes with varying impact. Population transcriptome and phenotype analyses highlighted the crucial role of accessible promoter regions as a genetic variation resource, influencing grain yield and quality in a core collection of wheat accessions. By integrating time-serial RNA-seq and ATAC-seq data, we constructed a hierarchical transcriptional regulatory network (TRN) governing starch and SSP biosynthesis, identifying 42 high-confidence novel candidates. These candidates exhibited overlap with genetic regions associated with grain size and quality traits, and their functional significance was validated through expression-phenotype association analysis among wheat accessions and TILLING mutants. In-depth functional analysis of *wheat abscisic acid insensitive 3-A1* (*TaABI3-A1*) with genome editing knock-out lines demonstrated its role in promoting SSP accumulation while repressing starch biosynthesis through transcriptional regulation. An elite haplotype of *TaABI3-A1* with higher grain weight was identified during the breeding process in China, and its superior trait was associated with altered *TaABI3-A1* expression levels. Additionally, we identified the potential upstream regulator, wheat GAGA-binding transcription factor 1 (TaGBP1), influencing *TaABI3-A1* expression. Our study provides novel and high-confidence regulators, presenting an effective strategy for understanding the regulation of SSP and starch biosynthesis and contributing to breeding enhancement.

## Introduction

Wheat, a major global staple, provides 20% of daily calorie intake and 15% of protein consumption worldwide. Given the growing population and changing climate, breeding efforts aim to enhance yield and quality (FAO., 2018). Endosperm development significantly influences grain yield and quality. Endosperm differentiation begins post-cellularization, with the outer layer forming the aleurone layer and inner cells becoming starch and protein storage cells (Liu et al., 2022). Grain filling involves the synthesis and accumulation of storage molecules like starch and gluten proteins (Nadaud et al., 2010). Starch constitutes 70% of dry weight, and seed storage protein (SSP) is pivotal for determining wheat flour’s end-use quality and imparting unique viscoelastic properties to dough (Kuchel et al., 2006; Xiao et al., 2022).

Wheat grain starch biosynthesis relies on a set of well-characterized enzymes and transporters, collectively known as starch synthesis-related genes (SSRGs). In cereals, 28 SSRGs, including ADP-glucose pyrophosphorylases (AGPases), granule-bound starch synthases (GBSSs), starch synthases (SSs), starch branching enzyme (SBE), debranching enzymes (DBEs), and starch/α-glucan phosphorylases (PHOs), play crucial roles in various steps of starch synthesis (Botticella et al., 2018). AGPase, the initial key regulatory enzyme, catalyzes the conversion of glucose-1-phosphate and ATP to ADPG. GBSS transfers ADP-glucose to linear glucose residues, forming amylose. SS elongates glucan chains, while SBE and DBE create the branched structure of amylopectin (Hannah and James, 2008; Jeon et al., 2010). Several transcription factors (TFs) have been identified as regulators of SSRGs in cereals. Rice Starch Regulator 1 (RSR1) negatively regulates starch synthesis-related enzyme-encoding genes (Liu et al., 2016). Basic leucine-zipper transcription factor 28 (TabZIP28) activates AGPase expression by binding its promoter, and its knockout leads to a 4% reduction in total starch content (Song et al., 2020). The endosperm-specific NAM/ATAF/CUC (NAC) TF TaNAC019 governs seed starch content and grain weight (Liu et al., 2020; Gao et al., 2021).

Gluten, a vital component in wheat influencing dough properties, comprises glutenins and gliadins (Wieser et al., 2023). Glutenins, including high-molecular-weight (HMW) and low-molecular-weight (LMW) subunits, form essential gluten polymers for dough elasticity (Lindsay and Skerritt, 1999). Gliadins, categorized as α/β-, γ-, ω-, and δ-gliadins, contribute to dough viscosity through interactions with glutenins (Urade et al., 2018). The intricate transcriptional regulation of gluten protein genes in developing wheat grain endosperm involves conserved *cis*-regulatory modules (CCRM1, CCRM2, CCRM3) enhancing HMW-GS expression (Ravel et al., 2014; Xiao et al., 2022). Segments CCRM1-1 and CCRM1-2 play pivotal roles in spatiotemporal gene expression. TFs are key regulators, where Storage Protein Activator (SPA) activates gluten gene expression, while SPA Heterodimerizing Protein (SHP) represses HMW-GS and LMW-GS genes (Boudet et al., 2019). TaFUSCA3, a B3-superfamily TF, activates HMW-GS subunit gene *Glu-1Bx7* through RY repeat binding (Moreno-Risueno et al., 2008). Prolamin box (P-box) binding factor (PBF), TaQM, and B3-domain TFs TaB3-2A1 regulate LMW-GS, gliadin, and *Glu-1* expression (Diaz et al., 2005; Dong et al., 2007; Zhu et al., 2018; Moehs et al., 2019; Xie et al., 2023). An R2R3 MYB TF, TaGAMyb, interacting with histone acetyltransferase General Control Nonderepressible (TaGCN5), activates HMW-GS via histone H3 acetylation (Guo et al., 2015). TaNAC019 interacts with TaGAMyb, activating *TaGlu* (Gao et al., 2021). NAC TF storage protein repressor (SPR), acts as a suppressor of SSP synthesis (Shen et al., 2021).

Genetic manipulation of starch and SSP genes is a prevalent strategy in wheat breeding. Notably, the *Wx* gene, crucial for amylose synthesis, has been extensively harnessed, enabling the creation of transgenic lines with diverse amylose content (AC) (Guzmán and Alvarez, 2016). The combination of null *wx* alleles facilitates the development of partial waxy and amylose-free waxy wheat (Guzmán et al., 2012). Induced mutations in *TaSBEII*, *TaSSII* and *TaSSIIIa* genes have yielded wheat varieties with elevated AC and resistant starch (Hazard et al., 2012; Hogg et al., 2017; Fahy et al., 2022). Effective enhancement of wheat quality has been achieved through the manipulation of main SSP genes, including *Grain protein content-B1* (*GPC-B1*) and *HMW-GS* alleles (Liang et al., 2010; Li et al., 2015; Tabbita et al., 2017). Beyond direct modification of starch and SSP genes, manipulating transcriptional regulators, such as *TaNAC019* and *TaNAC-A18*, has also been employed as a valuable tool in wheat breeding practices (Wang et al., 2023; Gao et al., 2021). For breaking the bottleneck of wheat yield and balancing the trade-off between yield and quality in wheat, it is necessary to systematically identify genetic regulators for wheat endosperm development, unravel the molecular mechanism of SSP and starch synthesis, and elucidate the interaction/coordination between these aspects.

Despite the identification of numerous genetic loci contributing to grain size, SSP, and starch-related traits through QTL analysis and genome-wide association studies (GWAS), only a few genes have been cloned (Rahimi et al., 2019; Pang et al., 2020; Khan et al., 2022; Li et al., 2022). This is attributed to inadequate mapping accuracy and limited insights into causal variants and genes due to linkage disequilibrium (Yu et al., 2014; Pang et al., 2020). To overcome these limitations, a comprehensive approach involving the generation of multi-layer data, including transcriptome (Rangan et al., 2017; Guan et al., 2022), translatome (Guo et al., 2023), metablome (Yin et al., 2024), and chromatin accessibility (Pei et al., 2023), has been implemented for the identification of key regulators in wheat spike and grain development (Lin et al., 2022; Luo et al., 2023).

In this study, we performed a thorough analysis of the transcriptome and epigenome profiles during the development of *cv.* Chinese Spring (CS) grains across various developmental stages. Our investigation delved into the dynamic transcriptome changes associated with starch and protein biosynthesis, connecting their expression variations with the phenotypic diversity within the population. Through the integration of transcriptional regulatory networks (TRN), GWAS data, TILLING mutant libraries, and population transcriptome information, we identified pivotal factors demonstrating the potential for synergistic regulation of SSP and starch synthesis.

## Results

### Transcriptome and chromatin landscapes during grain development in wheat

The endosperm of cereals offers the primary caloric and plant source-protein intake for the human diet (Liu et al., 2022). To understand the molecular basis of starch biosynthesis and SSP accumulation during endosperm development in wheat, we examined the transcriptional dynamics and chromatin landscapes of wheat endosperm during its major developmental stages, spanning from 0 to 22 days after pollination (DAP0-22) (Fig. 1, A and B). Due to the difficulty of dissecting, the embryo sac was harvested before DAP4, while the endosperm was isolated at DAP6 and later stages (Fig. 1, A and B). Regrettably, the high starch content during the late stages of endosperm development and the requirement of high purity of nuclei for ATAC-seq compared to CUT&Tag (Zhao et al., 2023), precluded the examination of chromatin accessibility status after DAP12 (Fig. 1B).

**Fig. 1.**
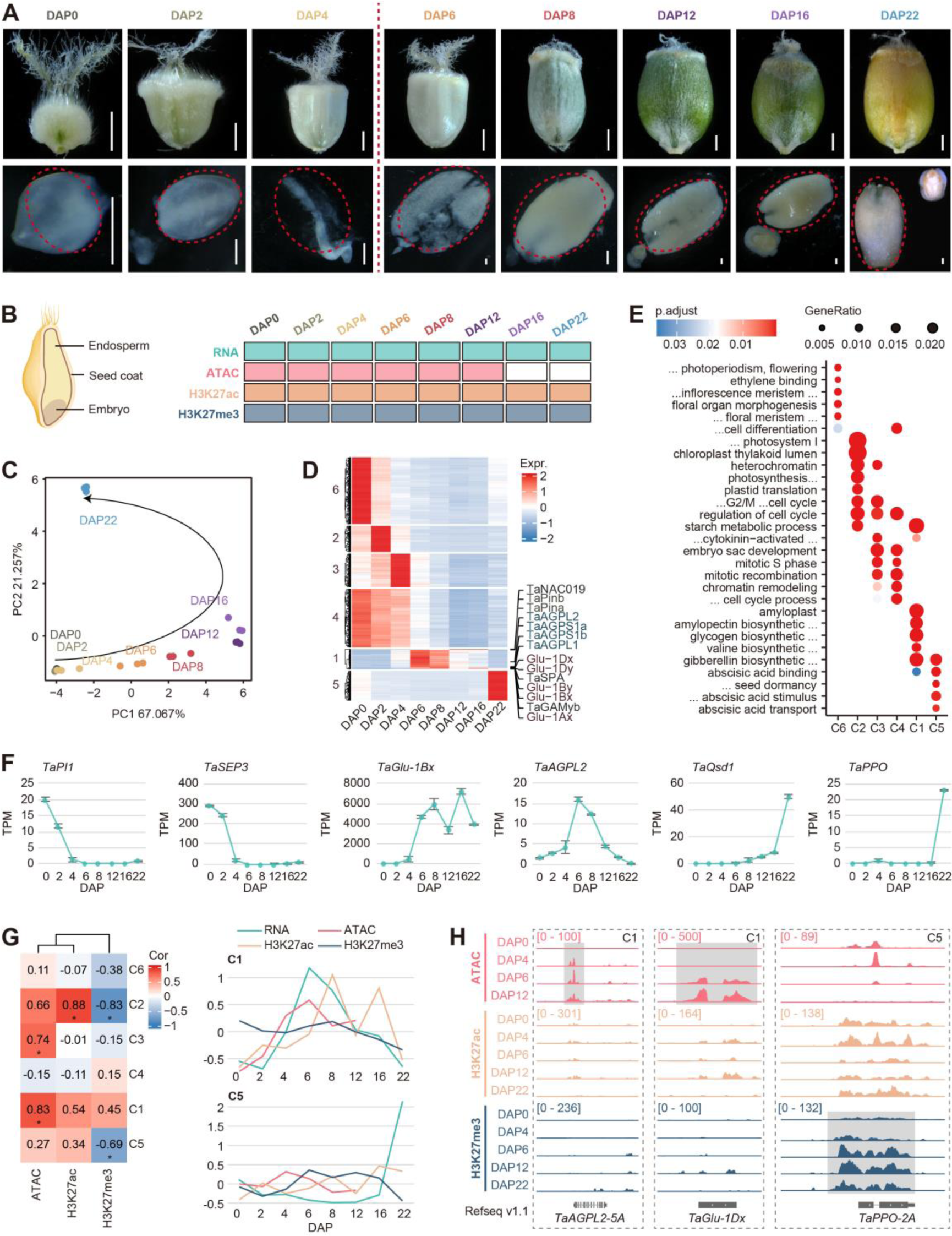
Transcription and epigenetic modification dynamics during grain development in wheat. **A.** Morphology of developing seeds from 0-22 days after pollination (DAP0-22) in wheat (upper); the dashed circle indicates tissue samples used for transcriptome and epigenome analysis at indicated DAPs (lower), Bar = 2 cm. **B.** Diagram indicating the general structure of wheat seed and experimental design of sampling. Endosperm tissues of different DAPs are used for RNA-seq, ATAC-seq and CUT&Tag analysis for H3K27ac, H3K27me3. **C.** PCA analysis of time-serial RNA-seq data, three bio-replicates were sequenced at each developing time point. **D.** K-mean clustering of differential expressed genes with grain development, starch and storage protein coding genes and some known transcription factors are highlighted. **E.** Go enrichment of different clusters in D. **F.** Representative genes from different clusters were exhibited with varied dynamic expression patterns during grain development from DAP0-22. **G.** Heatmap showing pearson correlation coefficient between genes expression and epigenetics modification in different clusters (left), the stars indicated the significant correlation with *P* ≤ 0.05. Line plot showing the dynamic of Z-score normalized average value of gene expression TPM and epigenetic modification peaks CPM at different DAP. **H.** IGV showing the various epigenetic modification tracks of representative genes *TaAGPL2* and *TaGlu-1Dx* from cluster 1 and *TaPPO-2A* from cluster5.

A total of 60,241 genes were found to be expressed in at least one sample during the development of the endosperm (table S1). Principal component analysis (PCA) revealed a progressive developmental trajectory, which could be grouped into three stages that mirror cellular transformations (Fig. 1C and fig. S1A). The global transcriptome was highly dynamic, with 96% of genes being considered as differential expressed genes (DEGs) among different samples (fig. S1B). Most triads among three sub-genomes were balanced expressed which was highly correlated with epigenetics modification levels (fig. S1, C to E). DEGs could be grouped into six clusters based on their temporal expression patterns, associated with different enrichments of GO and specific TF families (Fig. 1, D and E, and fig. S1F, and table S2). Interestingly, most genes involved in gluten protein and starch biosynthesis belonged to cluster 1, such as *TaAGPL2*, *TaGlu-1Bx*, which showed high expression level at DAP6 and DAP8, and extend expression till DAP16 (Fig. 1, D to F). For other clusters, Cluster 6 genes were predominantly expressed in unfertilized samples and were enriched with flower organ identity genes, such as *TaSEP3* (Schilling et al., 2020) and *TaPI1* (Paollacci et al., 2007) (Fig. 1, D to Lastly, Cluster 5 genes were specifically expressed at DAP22 and were related to seed dormancy, e.g. *TaQsd1* (Wei et al., 2019) and *TaPPO-2A* (Beecher and Skinner, 2011) (Fig. 1, D to F).

Next, we analyse the chromatin landscape dynamics during grain development. PCA of ATAC-seq data delineated a developmental trajectory, but not H3K27me3 and H3K27ac data (fig. S1G). Interestingly, different cluster of temporal expressed genes were correlated with distinct epigenetic modifications (Fig. 1, G and H, and fig. S1H). For instance, C5 cluster genes are negatively correlated with H3K27me3, whereas, C2 cluster genes showed positive correlation with H3K27ac and negative correlation with H3K27me3 dynamics. Notably, C3 and C1 cluster genes are both positive correlated with chromatin accessibility (Fig. 1, G and H, and fig. S1H). In particular, starch biosynthesis and gluten accumulation related genes *TaAGPL2-5*A and *TaGlu-1Dx* within C1 cluster show increased expression and chromatin accessibility from DAP6 to DAP12 (Fig. 1H). Whereas, *TaPPO-2A* within C5 cluster show specifically activation at DAP22, with the decline of H3K27me3 at the same time (Fig. 1H).

Thus, the dynamic high expression of various types of genes during endosperm development reflex the developing character of each stage. A distinct correlation between different cluster genes’ transcription and chromatin features suggests diverse regulation pattern of different stage-specific high expression genes.

### Expression and epigenetic modification of SSRGs, SSP genes and regulators during grain development

As starch and SSP biosynthesis and composition largely affects grain yield and quality in wheat (Kuchel et al., 2006), we further analyse the expression and epigenetic modification patterns of SSP coding genes (SSP), starch synthesis genes (Starch) and regulators (Reg) for various grain development traits including grain length (GL), grain width (GW), thousand grain weight (TGW), grain size (GS), SSP regulator (rSSP) during endosperm development (table S3). SSP-genes starting to express at DAP4 and decreasing after DAP20, while Starch-genes initiating at DAP4 and decreasing earlier after DAP8 (Fig. 2A). Reg-genes exhibited varied expression patterns, which fits with the different grain developmental traits associated (Fig. 2A) (Xiang et al., 2019; Liu et al., 2022). We analyzed the correlation between transcriptional dynamics with epigenetic modifications from DAP0 to DAP12 (Fig. 2, B and C, and fig. S2A), since ATAC-seq could be conducted only till DAP12 (Fig. 1B). For Starch-genes, transcription level shows high positive correlation with H3K27ac dynamics (R = 0.92), while to a less extend for chromatin accessibility (R = 0.78), especially after DAP8. Whereas, H3K27me3 dynamics is not well negatively correlated (R = −0.51) with transcription profiles. By contrast, for SSP-genes, chromatin accessibility and H3K27ac exhibits elegant positive correlation (R = 0.84, 0.73, respectively) while H3K27me3 shows negative correlation (R = −0.98) with transcription dynamics. The expression of Reg-genes in general are not associated with epigenetic modifications, may due to the diverse expression patterns of different regulators (Fig. 2, B and C, and fig. S2A). We further analyzed the subgenome expression preferences among triads of Starch- and Reg-genes. Whereas SSP-genes merely have homologs triads, we didn’t include it. The expression of Starch- and Reg-genes exhibited a primarily balanced pattern, while epigenetic modifications show greater subgenomic differentiation, indicating an integrated effect of various epigenetic modifications on transcription (Fig. 2D and fig. S2B).

**Fig. 2.**
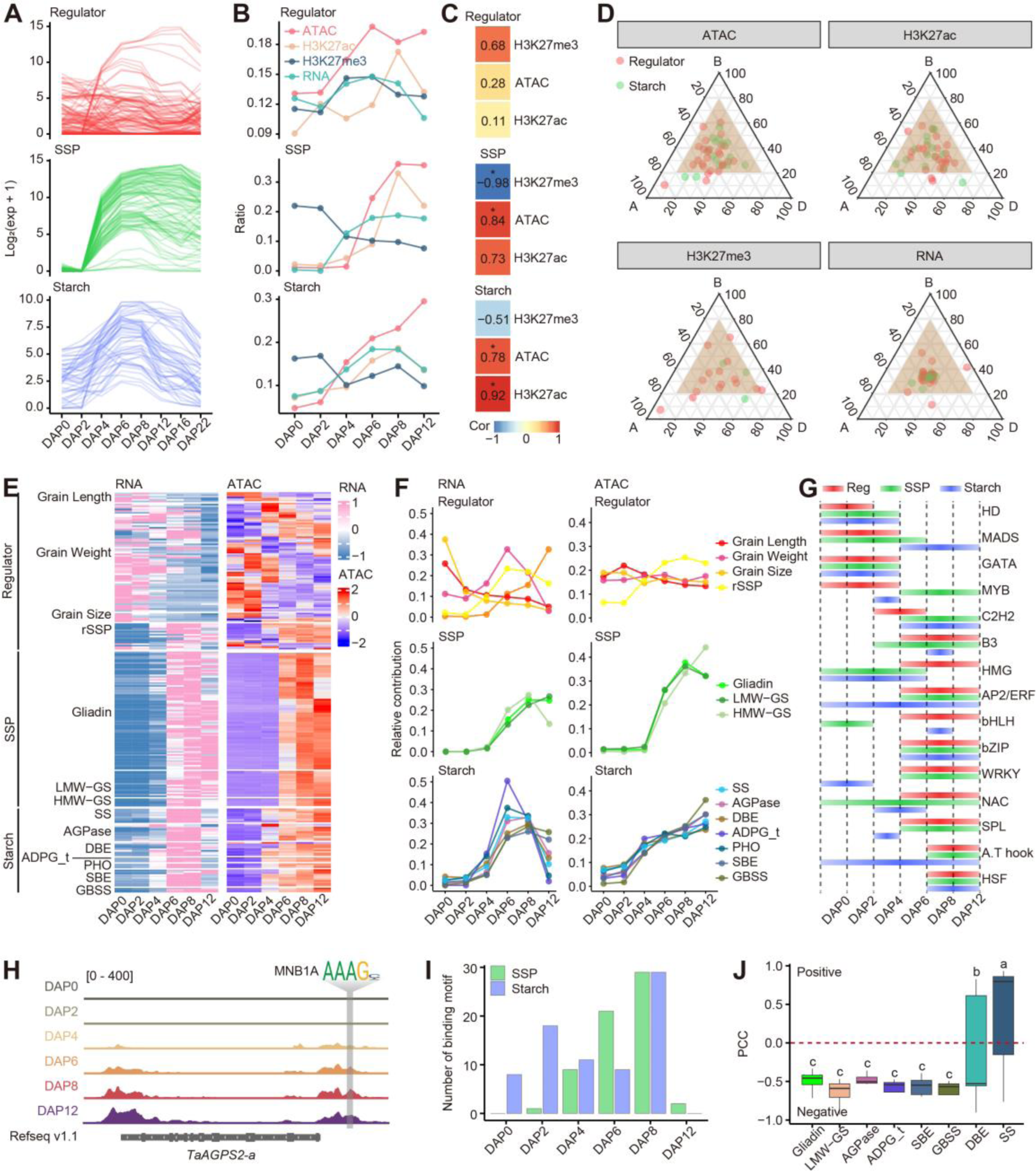
Robust correlation between epigenetic modification and dynamic expression of starch and SSP coding genes. **A.** The expression patterns of SSP coding, starch synthesis and regulators genes during DAP0-22. **B.** Correlation between genes’ expression and epigenetics modification in SSP-, Starch- and Reg-genes during DAP0-12. **C.** Heatmap showing Pearson correlation coefficient (PCC) between genes’ expression and epigenetics modification in three gene groups. The stars indicated the significant correlation with *P* ≤ 0.05. **D.** Ternary plot showing relative expression and epigenetics modification (ATAC, H3K27ac and H3K27me3) abundance of starch biosynthesis and regulators genes. Each circle represents a gene triad with A, B, and D coordinate consisting of the relative contribution of each homoeolog to the overall triad. Balanced triads are shown within brown shade. **E.** Gene expression (left) and chromatin accessibility (right) at proximal accessible regions (pACRs) of three-group genes and further sub-classification of each group. **F.** Expression (left) and chromatin accessibility at pACRs (right) patterns of each subgroup genes during endosperm development. The average TPM expression of sub-groups (based on Fig. 2E) at all stages was calculated. The relative contribute of each sub-group in each stage was then normalized to the average TPM expression of that sub-group divided by the total average TPM expression of all stages. **G.** Diagram shown the TF binding activity of each family accross developmental stages based on chromVAR. **H.** IGV showing the chromatin accessibility and binding motif of C2H2 family TF at the promoter of *TaAGPS2-a-7D*. Motif and footprint dynamic of C2H2 TF were shown. **I.** The number of NMNB1A binding footprint sites at starch and SSP coding genes during endosperm development. **J.** The boxplot of PCC between genes expression of each subgroup of starch and SSP coding genes and *NMNB1A*. The LSD multiple comparisons test was used to determine the significance of PCC differences among subtype genes. Different letters indicate a significant difference at *P* ≤ 0.05.

Notably, different sub-classes within the three gene sets exhibited relatively similar expression pattern associated with their biological function in mediating and regulating starch and SSP biosynthesis (Fig. 2, E and F). For instance, Reg-genes displayed two distinct expression patterns, with early-stage (DAP0-4) high expression related to GL and GS, whereas mid-stage (DAP6-12) high expression was associated with rSSP. Chromatin accessibility dynamics aligned well with the expression patterns of those sub-classes (Fig. 2, E and F). Similarly, within Starch-genes, *ADPG_t* expression initiated earliest, while *SBE* and *GBSS* expression occurred latest, associated with slightly different gains in chromatin accessibility. In contrast, all SSP-genes exhibited a similar expression and synchronized chromatin accessibility pattern (Fig. 2, E and F). In summary, chromatin accessibility showed a high correlation with sub-categories of Reg-, Starch-, and SSP-genes’ expression (fig. S2C).

Giving the strong correlation of chromatin accessibility with transcription dynamics, we employed chromVar to analyze the activity of TF binding motifs in the accessible regions of the three sets of genes (Fig. 2G). TF activities exhibited significant differences between early endosperm development (DAP0-4) and grain-filling stages (DAP6-12) (fig. S2D). C2H2, AP2, and bZIP displayed higher activity in the accessible promoter regions of SSP- and Starch-genes during grain-filling stages, indicating potential involvement in regulation of grain storage substance accumulation (Fig. 2G). For the HMW-GS, key factors regulating end-use quality, we discovered many conserved binding motifs of bZIP, NAC, and MYBs in the promoter region (fig. S2E). Notably, a gene belonging to the C2H2 TF family, Maize Dof1 (MNB1A) (Yanagisawa and Izui, 1993; Yanagisawa and Schmidt, 1999), exhibited strong activity in the accessible region of numerous SSP and starch biosynthesis genes during the DAP6-DAP12 stage (Fig. 2H and fig. S2F). The binding sites of MNB1A significantly increased during endosperm development, peaking at DAP8 (Fig. 2I). Notably, the expression of *MNB1A* was significantly negatively correlated with gliadin and LMW-GS coding genes as well as *AGPase*, *ADPG_t*, *SBE*, *GBSS*, while positively correlated with other starch synthesis coding genes like *SS*, *DBE*, and *PHOs* (Fig. 2J).

Thus, a robust correlation exist between epigenetic modifications and expression of starch biosynthesis and SSP-genes. This correlation is highlighted by the presence of specific cis-motifs and recognition TFs associated with accessible promoter regions in a timely manner.

### Expression variation of starch and SSP genes contributes to the diversity of grain size and quality within wheat population

Whether the variability in the expression of Starch- and SSP-genes contributes to the the final starch and storage protein contents, and the diversity observed in grain size and quality across wheat varieties? To address this, we assessed the coefficient of variation (CV) in the expression of Starch- and SSP-genes in 102 representative wheat varieties (Fig. 3A and fig. S3A, table S4). RNA sequencing of developing grains at DAP10 and DAP20 revealed a notably higher CV of SSP-genes, especially *ω-gliadins* varied the most, while *HMW-GS* showing the least variability. Among the Starch-genes, *ADPG_t* and *PHO* exhibited higher expression variations within the wheat population (Fig. 3A and fig. S3A). Interestingly, traits associated with protein content and SSP subunit composition displayed higher CV than traits related to starch content and pasting properties (fig. S3B).

**Fig. 3.**
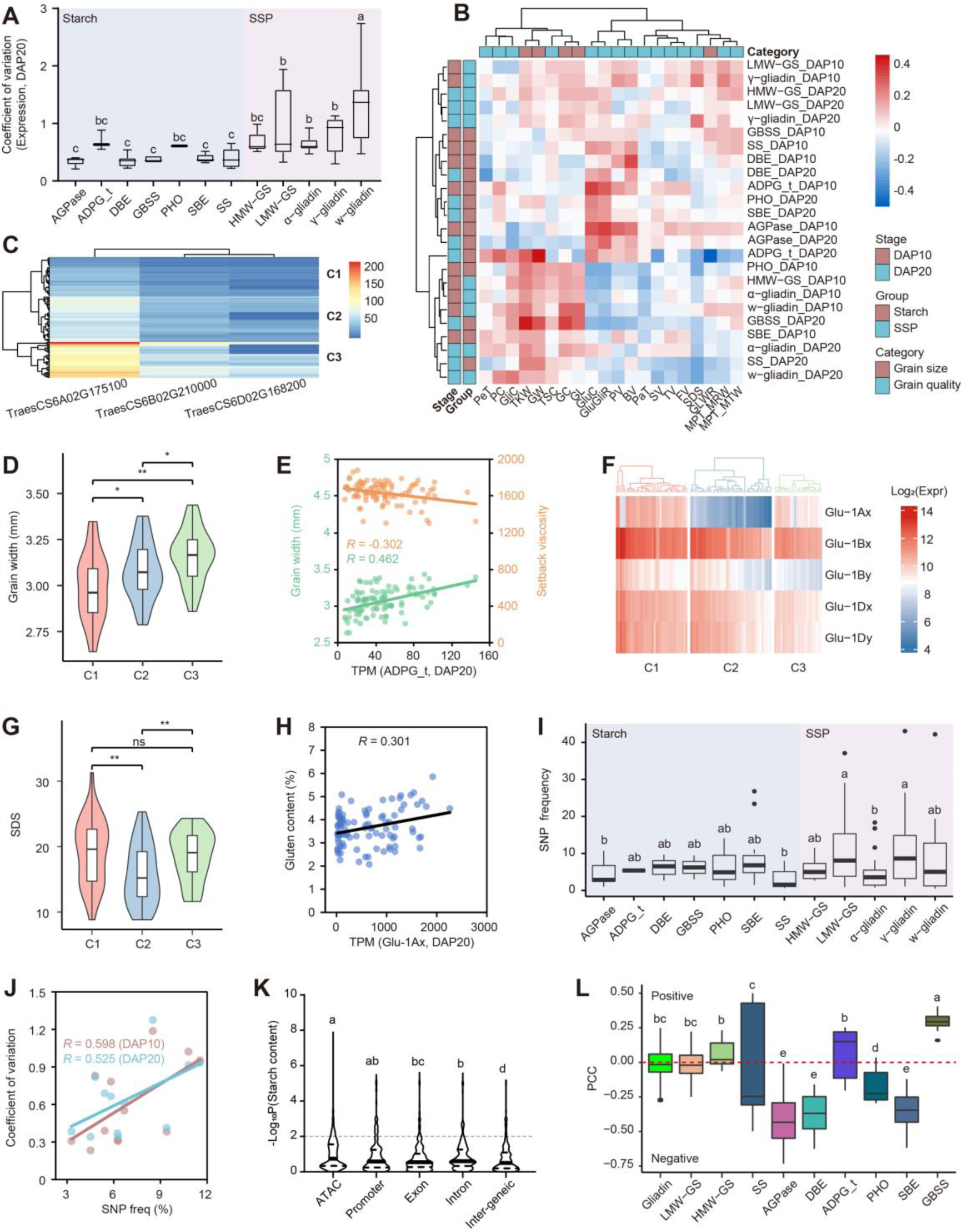
Variation in the regulatory region alters expression and influences grain size and quality of wheat population. **A.** The comparison of coefficient of variation (CV) in expression levels for each subtype genes within the core collection of 102 wheat accession at DAP20. The box denote the 25^th^, median and 75th percentiles, and whiskers indicate 1.5× interquartile range. The LSD multiple comparisons test was used to determine the significance of CV differences among subtype genes. Different letters indicate a significant difference at *P* ≤ 0.05. **B.** A heatmap showing the correlation between phenotypic traits and the expression level of subtype genes. **C.** Expression level of *ADPG_t* genes in developing grains at DAP20 within the wheat population. C1, C2, C3 areclustered by the TPM value with “complete” method using pheatmap. **D.** Comparison of the grain width of wheat accessions among three groups. The student’s t-test was used to determine the statistical significance between two groups. *, *P* ≤ 0.05; **, *P* ≤ 0.01. **E.** Scatter plot of grain width (green dots) and setback viscosity (brown dots) against the expression level values of *ADPG_t* in corresponding accession at DAP20. Each dot denotes one accession, and the lines represent the regression trend calculated by the general linear model. **F.** Expression level of *HMW-GS* genes in developing grains at DAP20 within wheat population. C1, C2, C3 are clustered by the Log_2_(TPM value) with “complete” method using pheatmap. **G.** Comparison of the SDS value of wheat accessions among three groups. The student’s t-test was used to determine the statistical significance between two groups. *, *P* ≤ 0.05; **, *P* ≤ 0.01; ns, no significant difference. **H.** Scatter plot of gluten content against the expression level values of *Glu-1Ax* in corresponding accessions at DAP20. Each dot denotes one accession, and the line represents the regression trend calculated by the general linear model. **I.** Comparison of the SNP frequency in open chromatin regions among each subtype genes. The SNP frequency was calculated using SNP number in the Watkins population divided by the length of ATAC-seq peaks. Box denote the 25^th^, median and 75th percentiles, and whiskers indicate 1.5×interquartile range. The LSD multiple comparisons test was used to determine the significance of CV differences among subtype genes. Different letters indicate a significant difference at *P* ≤ 0.05. **J.** Scatter plot of the expression CV at DAP10 (brown dots) and DAP20 (cyan dots) against the SNP frequency of each subtype of genes. Each dot denotes one subtype, and the gray line represents the regression trend calculated by the general linear model. **K.** Comparison of the −log_10_(*p* value) of SNPs with starch content among different genomic features. The LSD multiple comparisons test was used to determine the significance differences among subtype genes. Different letters indicate a significant difference at *P* ≤ 0.05. **L.** The Pearson correlation coefficient (PCC) of expression level in population between *MNB1A* and SSP and starch synthesis genes. Box denote the 25^th^, median and 75th percentiles, and whiskers indicate 1.5×interquartile range. The LSD multiple comparisons test was used to determine the significance of PCC differences among subtype genes. Different letters indicate a significant difference at *P* ≤ 0.05.

We further examined the relationship between the expression levels of different subtypes of Starch- and SSP-genes, with grain size and quality traits. Starch-genes, more active in the early stages of endosperm development, showed a stronger correlation with starch content and pasting properties at DAP10 than at DAP20 (Fig. 3B and table S5). Conversely, SSP coding genes exhibited a higher correlation with traits related to protein content and SSP subunit composition at DAP20 (Fig. 3B). Among the 102 wheat varieties, *ADPG_t* expression varied significantly, with the highest in C3, followed by C2, and the lowest in C1 (Fig. 3C). Remarkably, the GW of accessions in each cluster followed the order of expression level (C3 > C2 > C1), and setback viscosity (SV) exhibited the opposite trend (Fig. 3D and fig. S3C). Correspondingly, *ADPG_t* expression levels were significantly positively correlated with GW and negatively with SV (Fig. 3E). Similarly, the expression level of *TaGlu-1*, especially *TaGlu-1Ax*, showed a significant positive correlation with gluten content and SDS value in the population (Fig. 3, F to H, and fig. S3, D and E). These findings suggest that the expression variation of SSP and starch synthesis genes may contribute to the observed diversity in grain size and quality within the wheat population.

Considering the crucial role of open chromatin regions in regulating genes related to SSP and starch synthesis, we explored whether natural variations in these regions contribute to expression variations. The SNP frequency in open chromatin regions of SSP genes, particularly *LMW-GS* and *γ-gliadins*, was higher than that of starch synthesis genes, while *AGPS* and *SS* exhibited lower SNP frequencies (Fig. 3I). Consistently, this SNP frequency in open chromatin regions strongly correlated (R > 0.5) with their expression variability among cultivars in the population, both in developing grains at DAP10 and DAP20 (Fig. 3J). Additionally, natural variations in open chromatin regions had higher detection power, contributing to phenotypic variation within the population (Fig. 3K and fig. S3F and G). A larger proportion of SNPs were significantly associated with starch and protein content in a previously characterized wheat population (Cheng et al., 2023), compared to other genomic regions (Fig. 3K and fig. S3, F and Notably, the expression of MNB1A, which binds to the SSP and starch synthesis genes, is significantly correlated with starch synthesis genes within the population, albeit with varying relations among different subtypes (Fig. 3L). Collectively, natural variations in the open chromatin regions of SSP and starch synthesis genes may represent causal variations contributing to their expression difference and the observed diversity in grain size and quality within the wheat population.

### Transcriptional regulatory networks reveals the regulation principle of storage production accumulation

Revealing the intricate regulatory mechanisms governing starch biosynthesis and SSP accumulation during endosperm development is crucial for understanding storage production accumulation. To achieve this, we constructed hierarchical transcriptional regulatory networks (TRNs) by integrating gene co-expression information (GENIE3) and ATAC-seq footprint data (fig. S4A), resulting in a complex TRN with 3,187,686 connections and 60,973 nodes involving 1,856 transcription factors (TFs) (Fig. 4A). TFs of the same family exhibited similar binding motifs and shared common targets, revealing their coordinated regulation (Fig. 4A). Notably, known TFs implicated in wheat end-use quality, such as TaERF3, TaFUSCA3, TaSPR, TaARF25, TaNACs, and TaGPC-1 and NAC TFs were identified in the TRNs (Fig. 4A).

**Fig. 4.**
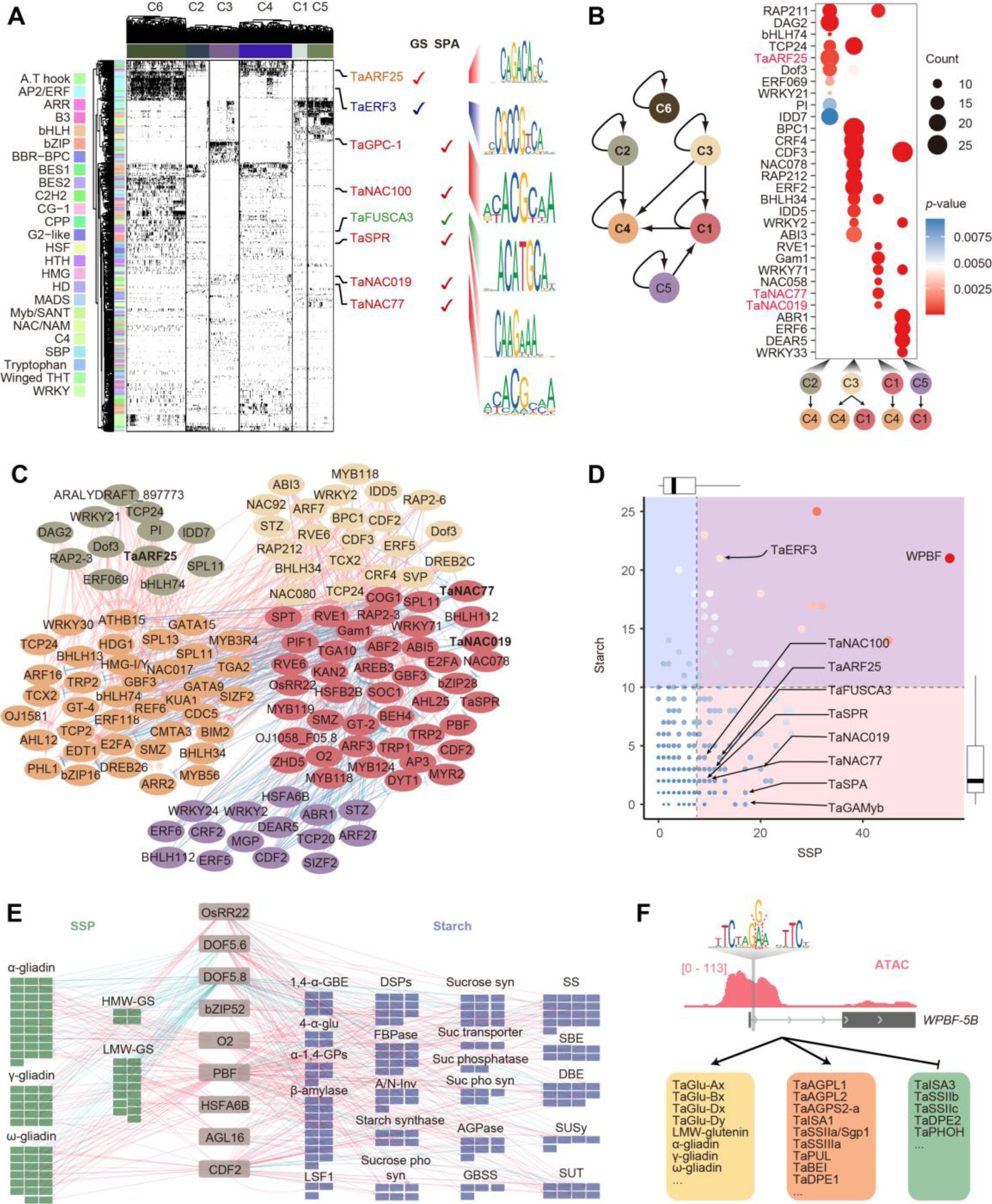
Transcriptional regulatory networks (TRNs) during endosperm development. **A.** The two dimensional transcriptional regulatory matrix for TFs-targets. Each column represented a target gene and each row represented a TF. The target genes were ordered by clustering result in Fig. 1d. The black color represents a regulation relationship between TFs and targets, while the white color means no regulation. Known TFs regulating grain size (GS) and storage production accumulation (SPA) were highlighted. The binding motifs of representative TFs were exhibited. **B.** Transcriptional regulatory relationship between different cluster genes (left) and the enrichment of conducting TFs (right). **C.** TFs-TFs regulatory networks among clusters C1, C2, C3, C4, C5. Enriched TFs were shown in color circles. The color code for each cluster was the same as shown in B. Known TFs were indicated. **D.** Different TFs could directly regulate multiple SSP (x-axis) and starch (y-axis) biosynthesis related genes. TFs with more than ten SSP and/or starch biosynthesis genes as targets were highlight in colored shadow. Known TFs were indicated. The outer boxplot represented the statistics of the number of SSP and starch genes. **E.** Co-regulation of starch biosynthesis and SSP composition related genes by common TFs. SSP and starch biosynthesis genes were colored in red and blue, respectively. Each function pathway genes were clustered. The red line between TFs and target represented the positive regulation, while blue line represented the negative regulation. **F.** Regulation of *WPBF* on starch biosynthesis and SSP composition related genes. The proximal accessible region of *WPBF* included an SNP in a TF binding motif. WPBF could positively regulate SSP or starch biosynthesis genes, as shown by arrow, and negatively regulate starch biosynthesis genes, shown by break line.

Furthermore, the TRNs highlighted the conservation of transcription factor binding sites (TFBS) across the different clusters, emphasizing the intricate regulatory relationships governing gene expression during endosperm development (fig. S4, B and C). Notably, genes within cluster 1, enriched in protein-biosynthesis, exhibited regulation by clusters 3 and 5, suggesting coordinated control of key processes (Fig. 4, B and C). The specific enrichment patterns like TaARF25 and NAC TFs for the regulation of cluster 4 genes by clusters 2 and 1, respectively, underscored the nuanced regulatory networks orchestrating endosperm development (Fig. 4B). Examining the hierarchical positioning of established TFs associated with storage production, including WPBF, TaSPR, TaSPA, TaGAMyb, TaNAC019, and TaNAC77, provided valuable insights into their direct influence on storage production accumulation (Fig. 4C). In addition to several documented TFs known to influence storage production, we identified over a hundred TFs, which could exert positive or negative regulation on either or both of Starch- and SSP-genes (Fig. 4, D and E and fig. S4D). Among those, majority TFs show negative regulation for both SSP and starch biosynthesis genes, whereas, less number of TFs can simultaneously regulate Starch- and SSP-genes positively. In addition, several TFs could regulate the expression of Starch- and SSP-genes in a an opposite way, such as WPBF, ERFs, MNB1A, ABI3, suggesting their potential function in balancing SSP and starch biosynthesis (fig. S4D).

Notably, WPBF emerged as a central player in this regulatory network, displaying a remarkable capacity to simultaneously positively and negatively regulate a multitude of storage production genes (Fig. 4, D and E). For instance, WPBF may positively regulate SSP such as HMW-GS, LMW-GS and glidins genes, positively regulate certain starch biosynthesis genes, such *TaAGPase*, *TaSA*, *TaBEI* while negatively regulate others, such as *TaISA3, TaSSII, TaDPE2* and *TaPHOH*. The identification of a SNP at the binding site of WPBF within the wheat population added an additional layer of significance, suggesting potential variations in its direct regulatory impact (Fig. 4F).

Therefore, the generated TRN enhances understanding of transcriptional regulation of starch biosynthesis and SSP accumulation. Relationships among TFs, binding motifs, and genetic variations within the population offer insights for optimizing storage production.

### Identification of novel regulators governing grain yield and quality in wheat

Based on the constructed TRN, a substantial portion of known TFs involved in the structural organization and direct regulation of storage production biosynthesis genes was identified (Fig. 4, C and D). The integration of these two gene lists yielded a total of 436 core TFs (Fig. 5A and table S6). Among them, 26 TFs have been functionally validated in wheat for their role in regulating grain size or seed quality, including factors like TaGPC-1 (Uauy et al., 2006), TaNAC019 (Liu et al., 2020; Gao et al., 2021), TaSPR (Shen et al., 2021). Additionally, 15 TFs are orthologous to regulators for kernel development in rice, such as OsbZIP60 (Cao et al., 2022; Yang et al., 2022), OsDOF17 (Wu et al., 2022) and OsAPG (Heang and Sassa, 2012). Notably, the majority of the TFs, comprising 395 candidates, are novel and have unknown functions in wheat, with enrichment in ERF, NAC, ARF, and B3 TF families (table S6).

**Figure 5.**
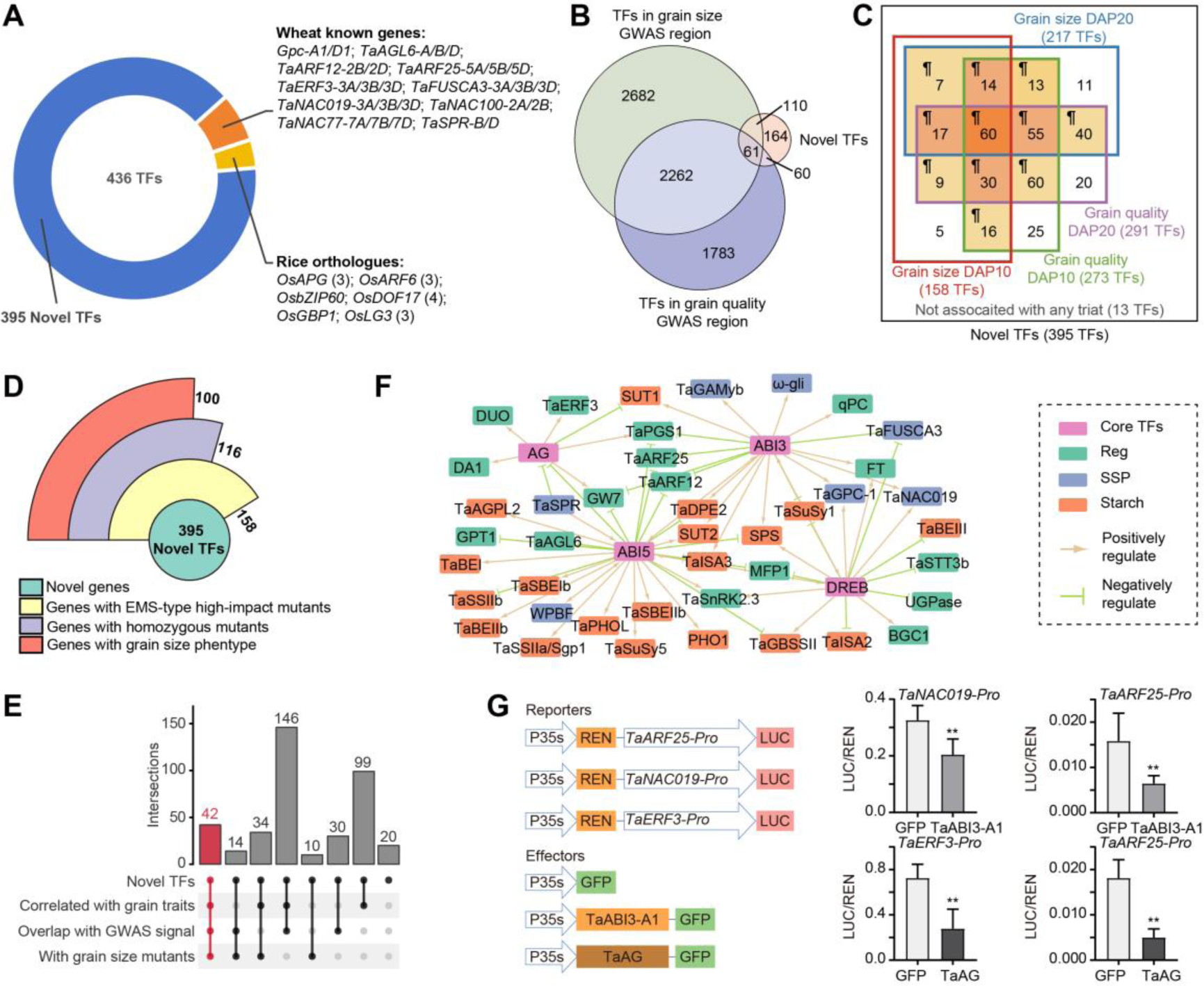
Filter and validation of potential key TFs in shaping grain development traits. **A.** Characteristic of core TFs from TRN analysis. **B.** Venn-diagram showing the overlapping of novel TFs identified from TRN with TFs located in the interval of significant GWAS signal for grain size and grain quality. **C.** Summary of novel TFs identified from TRN showing association between TFs’ expression with grain size and quality traits variation in a core collection of wheat accessions. The genes identified in more than two groups were marked with background color, with genes overlapped in more groups in darker color. ¶, the genes reserved for further analysis. **D.** Summary of novel TFs with different categories of KN9204 TILLING mutant lines. **E.** Overlapping of the novel TFs supported by GWAS signal, expression-phenotype correlation and morphological defects of TILLING mutants. **F.** A case show of TRN module containing core TFs such as TaABI3, TaABI5, TaDREB, TaAG and the potential targets involved in SSP and starch biosynthesis, as well as the known regulators. Color codes are as indicated. **G.** Dual luciferase reporter assays showing the transcriptional regulation of TaABI3, TaAG to potential targets. Student’s t-test was used for the statistical significance. *, *P* ≤ 0.05; **, *P* ≤ 0.01.

To assess the efficacy of identifying novel regulators through TRN, we conducted a comprehensive evaluation in three aspects. Firstly, we combined available GWAS data for grain size and quality (table S7) with the 395 novel candidates, resulting 231 TFs within 3 Mb regions centered around GWAS signals, with 171 and 121 TFs for grain size and grain quality related traits, respectively (Fig. 5B and table S8). Secondly, we correlated the expression levels of 395 novel candidate genes at DAP10 and DAP20 with grain size and quality-related traits in the 102 representative wheat varieties. For 158 TFs, the expression level at DAP10 were significantly correlated with at least one grain size related traits (GS-DAP10 group). Similarly, 273, 217 and 291 TFs were found in the GQ-DAP10 group, GS-DAP20 group and GQ-DAP20 group, respectively (Fig. 5C and table S8 and S9). In total, we identified 321 TFs showing significant correlation in more than two groups (Fig. 5C and table S9). Thirdly, we conducted a robust investigation of grain size changes in gene-indexed EMS mutant lines in hexaploid wheat *cv.* KN9204 (Wang et al., 2023a). Of the 395 novel TFs, 158 TFs were found to have at least one mutant line that containing loss-of-function mutation, and homozygous mutant lines of 100 out of 116 TFs (86.21%) exhibited altered grain size or morphology (Fig. 5D and table S10). Collectively, a substantial portion of the identified novel TFs exhibited associations with grain-related traits across genome variation, transcriptional divergence, and protein function levels.

By integrating these verification strategies, 42 novel TFs supported by all approaches were identified for in-depth study (Fig. 5E and table S8). Focusing on the TRN for these 42 key TFs, a transcription regulatory module involving TaABI3, TaAG, TaDREB and TaABI5 was revealed, synergistically regulating grain size regulators (e.g., GW7 and TaARF12), starch biosynthesis genes (e.g., SUT1, TaDPE2, and SPS), and storage protein formation (e.g., TaFUSCA3, TaGPC-1 and TaNAC019) (Fig. 5F). We further validated some of the transcriptional regulatory modules by luciferase reporter assay in tobacco leaves, such as the inhibition of TaABI3 on *TaNAC019* and *TaARF25*, and the repression of TaAG on *TaERF3* and *TaARF25* (Fig. 5G). These findings underscore the accuracy and reliability of predicted grain development regulators by the TRN.

### TaABI3-A1 negatively modulates starch biosynthesis while activating storage proteins accumulation

To delve into the functional insights of key potential factors identified within the TRN in wheat endosperm development, we conducted an in-depth investigation of a novel regulator, TaABI3-A1 (*TraesCS3A02G417300*). Through *in situ* hybridization, *TaABI3-A1* displayed distinctive expression patterns during grain development, with higher expression level particularly in the endosperm and aleurone layer, but lower expression levels in the seed coat at DAP6 (Fig. 6A). To elucidate TaABI3’s role in wheat grain development, we generated two knockout lines with premature stops in all three homoeologues using CRISPR/Cas9 in wheat *cv.* JM5265 (Fig. 6B and fig. S5A).

**Figure 6.**
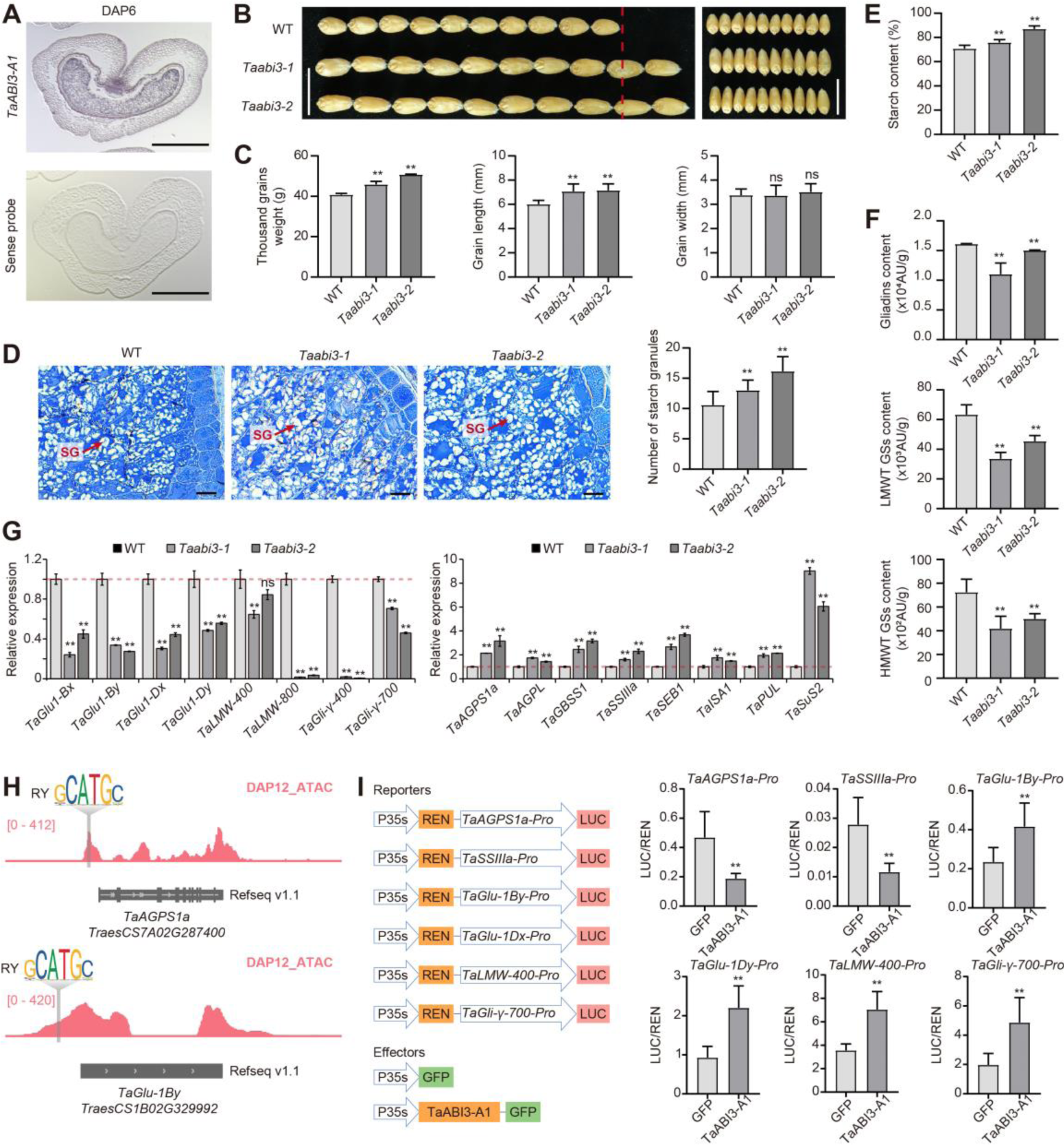
TaABI3-A1 regulates the grain size and quality of wheat. **A.** Spatiotemporal expression pattern of *TaABI3-A1* in DAP6 grain as indicated by *in situ* hybridization. Sense probe served as negative control. Scale bars = 100 μm. **B.** Loss-of-function *Taabi3* mutants significantly increases the grain size in wheat. Red line indicated the length of ten wild-type seeds. **C.** Quantification of grain size related traits between the wild-type plants and *Taabi3* mutant lines. Student’s t-test was used to determine the difference significance between *Taabi3* mutant and wild-type. *, *P* ≤ 0.05; **, *P* ≤ 0.01; ns, no significant difference. **D.** Representative cross sections of DAP30 grains for wild-type and *Taabi3* mutant lines (Left). Red arrows indicate the starch granules (SG). The quantification of SG numbers was shown in bar graph (Right). Student’s t-test was used to determine the difference significance between *Taabi3* mutant and wild-type. *, *P* ≤ 0.05; **, *P* ≤ 0.01. **E, F.** Comparison of the content of starch (E) and SSP components (F) between wild-type and *Taabi3* mutant lines. Three repeats were carried out for each sample to measure the amounts of starch, HMW-GSs, LMW-GSs, and gliadins content. **G.** The relative expression level of SSRGs and SSP-related genes in wild-type and *Taabi3* mutant lines. RT-qPCR data were normalized to *TaACTIN*, with values from three replicates shown as mean ±sd. Statistical significance was determined by Student’s t-test. *, *P* ≤ 0.05; **, *P* ≤ 0.01; ns, no significant difference. **H.** IGV screenshot showing the accessible chromatin regions of *TaAGPS1a*, and *TaGlu-1By* with a gray vertical line indicating the RY motif. **I.** Dual luciferase reporter assays showing the transcriptional regulation of TaABI3-A1 on SSRGs and SSP-related genes. Schematic diagram showing the vectors used. The relative value of LUC/REN was indicated. Relative LUC/REN values from six replicates shown as mean ± sd. Statistical significance was determined by Student’s t-test. *, *P* ≤ 0.05; **, *P* ≤ 0.01.

Field growth evaluations of grain size phenotypes for *Taabi3-a/b/d*, alongside the wild-type JM5265, revealed a significant increase in GL and TGW in *Taabi3-a/b/d* compared to the wild-type (Fig. 6C). However, *TaABI3* knockout did not alter grain width (Fig. 6C). To understand the underlying mechanisms, we examined endosperm cells in developing seeds at DAP15. Coomassie brilliant blue staining revealed fewer starch granules in *Taabi3* endosperm cells compared to the wild-type (Fig. 6D) and increased total starch content in mature seeds (Fig. 6E). Additionally, reverse-phase high-performance liquid chromatography (RP-HPLC) indicated reduced levels of HMW-GS, LMW-GS, and gliadins in *Taabi3* mutants (Fig. 6F), highlighting TaABI3’s role in regulating SSP. Consistence with the grain developmental defects, in the *Taabi3* mutants, the expression level of SSP-related genes were significantly reduced, among which *TaGli-400* and *TaLMW-800* drastically declined by dozens of times (Fig. 6G). By contrast, some key genes controlling sugar or starch synthesis were up regulated in the *Taabi3* mutants, compared with wild-type (Fig. 6G). Notably, the Wheat sucrose synthase 2 (*TaSUS2*), significantly associated with TGW in wheat (Jiang et al., 2011), was more than six times higher in *Taabi3* mutants than wild-type (Fig. 6G). To explore the regulatory mechanism, we utilized ATAC-seq data to identify the RY motif recognition by ABI3 TF family in the promoters of these starch and SSP biosynthesis related genes (Fig. 6H). Furthermore, a dual luciferase transcriptional activity assay revealed a positive transcriptional regulatory of TaABI3-A1 on SSP-related genes *TaGlu-1By*, *TaGlu-1Dx*, *TaLMW-400*, and *TaGli-γ-700*, while suppressed the expression of *TaAGPS1a* and *TaSSIIIa,* in tobacco leaves (Fig. 6I and fig. S5B).

In summary, TaABI3-A1 negatively regulates the grain size, primarily through the transcriptional inhibition of starch biosynthesis, while promoting storage protein accumulation by activating SSP-related genes.

### Genetic variation of *TaABI3-A1* is associate with grain weight and quality

To explore the link between genetic variation in *TaABI3-A1* and grain-related traits in wheat, we examined allelic diversity in the Chinese wheat mini-core collection (MCC), a panel of 287 representative varieties (Li et al., 2022). In the *TaABI3-A1* genomic region, 15 polymorphic sites, including six SNPs in the promoter, five SNPs on the exon, and two SNPs with two 1-bp deletions at intron, formed three haplotypes (Fig. 7A). Hap-1, represented by 42 accessions, showed significantly higher grain length, width, and weight than Hap-2 and Hap-3 (Fig. 7B and table S11). Statistical analysis revealed that Hap-1 had superior grain size compared to Hap-2 and Hap-3, designating it as the excellent haplotype.

**Fig. 7.**
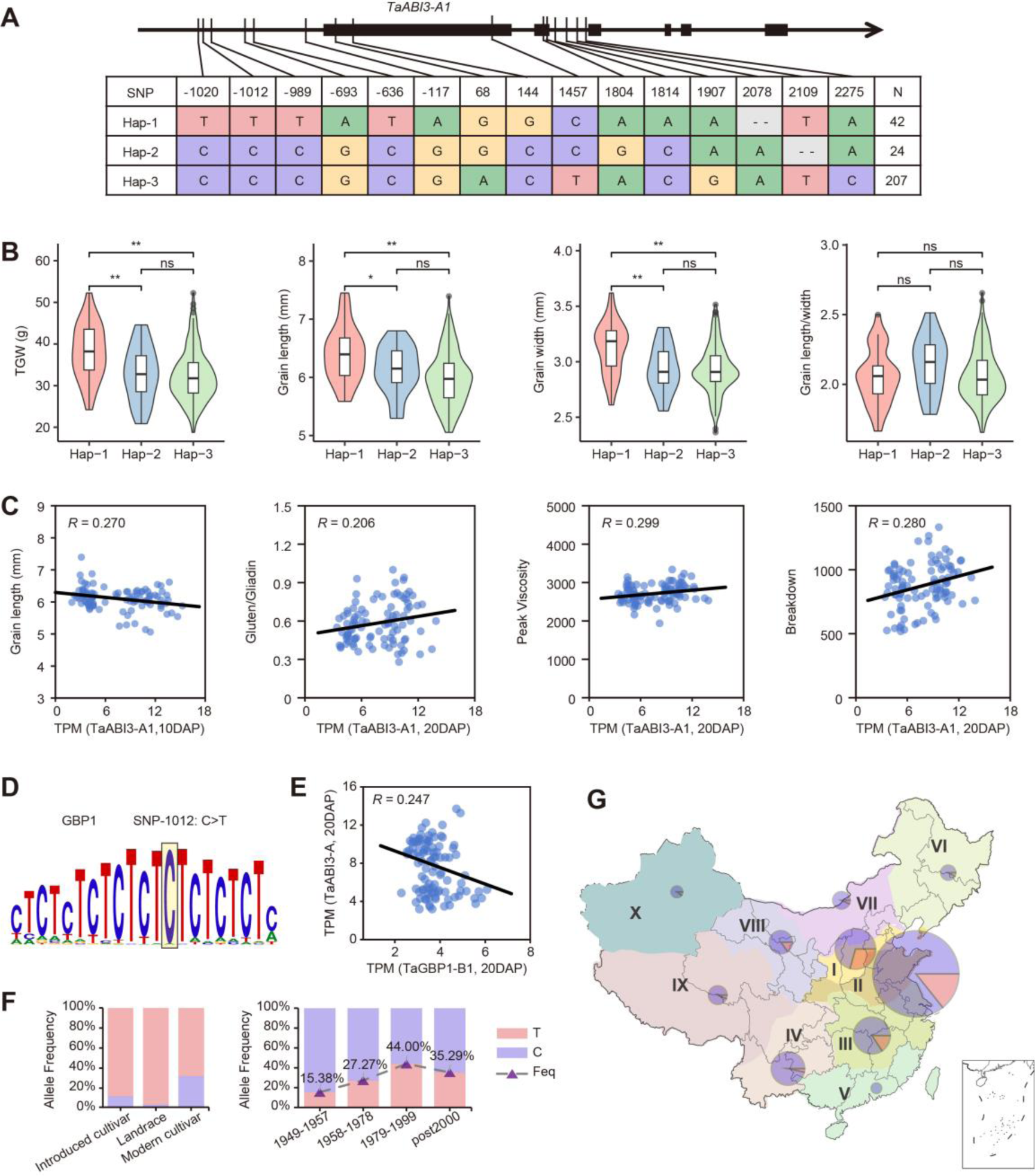
Haplotype analysis of *TaABI3-A1* and breeding selection of elite allele. **A.** Schematic diagram showing the polymorphism for each haplotype of *TaABI3-A1* in the Chinese wheat mini-core collection (MCC) wheat collection. The coordinate is related to transcription start site (TSS). **B.** Violin plot indicating the comparison of grain size related traits among wheat accession with different haplotypes of *TaABI3-A1*. The student’s t-test was used to determine the statistical significance between two groups. *, *P* ≤ 0.05; **, *P* ≤ 0.01; ns, no significant difference. **C.** Scatter plot showing the correlation between grain length, peak viscosity, breakdown value and gluten:gliadin ratio with the expression of *TaABI3-A1*. Each dot denotes one accession, and the black line represents the regression trend calculated by the general linear model. **D.** The SNP-1012 located in the accessible chromatin region of *TaABI3-A1* promoter under the core DNA-binding motif of TaGBP1. **E.** The correlation between the expression level of *TaGBP-B1* and *TaABI3-A1* in the developing grains of a core collection of 102 wheat accessions (DAP20). Each dot denotes one accession, and the black line represents the regression trend calculated by the general linear model. **F.** The percentages of accessions carrying different allele of SNP-1012 in each categories (left) and during the different breeding process in China (right). **G.** The percentage of accessions carrying different allele of SNP-1012 in each ecological zones of China. The size of pie charts in the geographical map shows the number of accessions, with percentages of the two allele in different colors (C-allele, purple; T-allele, pink).

To assess if *TaABI3-A1* expression contributes to grain size and quality variations, given its negative regulation of grain length and starch content (Fig. 6, A to D), we examined its expression at DAP10. A significant negative correlation between grain length and *TaABI3-A1* expression was observed, and further flour quality analysis showed a positive correlation with peak viscosity, breakdown value, and gluten: gliadin ratio (Fig. 7C). These results suggest that *TaABI3-A1* expression contributes to variations in grain size and quality among wheat varieties, impacting different traits to varying extent.

Causal variations influencing *TaABI3-A1* expression and superior traits were considered to be SNPs in the promoter region, as these sequence differences between Hap-1 and the other haplotypes primarily resided in the promoter region. Among the six Hap-1 promoter specific SNPs, SNP-1012 (C>T) was identified at core binding motifs and predicted to disrupt the binding of TaGBP1 at the *TaABI3-A1* promoter (Fig. 7D). The rice ortholog OsGBP1 is known to negatively regulate kernel length and weight (Gong et al., 2018). Correspondingly, the expression level of *TaABI3-A1* was negatively correlated with that of *TaGBP1-B1* in DAP20 grains (Fig. 7E). Thus, the T-to-C variation in promoter may increases *TaABI3-A1* expression by reducing TaGBP1 repression, associating with superior grain weight and quality in wheat.

To assess whether the excellent haplotype was selected in China’s breeding process, we analyzed SNP-1012 allele frequency in the MCC population. The percentage of accessions with the T-allele was notably higher in modern cultivars than in landraces and introduced cultivars from other countries (Fig. 7F). Interestingly, the T-allele frequency steadily increased from 15.38% in 1949-1957 to 44.00% in 1978-1999, slightly decreasing in post-2000 cultivars. Additionally, distinct allele distribution was observed in major Chinese agro-ecological zones, with zones I, II, and III showing a higher T-allele frequency than other zones (Fig. 7G). Collectively, the bigger grain T-allele of *TaABI3-A1* was selected during the breeding process of China.

In summary, natural variations in SNP-1012 in the *TaABI3-A1* promoter associate with its expression level and superior grain weight in wheat, undergoing selection in China’s breeding process.

## Discussion

Understanding the molecular mechanisms governing starch biosynthesis and SSP accumulation is vital for breeding improved wheat varieties (Muqaddasi et al., 2020; Xiao et al., 2022). Efforts to identify enzymes for starch and SSP, along with genetic variations associated with them, have linked to enhanced grain quality (Liang et al., 2010; Li et al., 2015; Tabbita et al., 2017). However, a gap exists due to the lack of systematic transcriptional regulation analysis of starch and SSP coding genes. In this study, we generated transcriptome and epigenetic profiles across key endosperm development stages (Fig. 1). Our approach identified specific open chromatin regions regulating starch and SSP genes (Fig. 2). By integrating expression data and *cis*-*trans* regulation, we built a hierarchical transcriptional regulatory network (TRN) for starch and SSP (Fig. 4), efficiently identifying novel regulators for grain yield and quality (Fig. 5).

### Significance of regulatory regions in shaping starch and SSP diversity

Various steps are essential for producing starch and different gluten proteins with distinct characteristics, and previous efforts have identified enzymes and non-enzymatic factors involved in this process (Vasil and Anderson, 1997; Morell et al., 2001; Delcour et al., 2012). Studies have explored coding variations in these genes contributing to diverse starch and SSP products (Gil-Humanes et al., 2011; Wang et al., 2020b). Our research reveals that dynamic epigenetic features, particularly chromatin accessibility, H3K27ac, and H3K27me3, collectively regulate SSP genes, with a less pronounced effect of H3K27me3 on starch biosynthesis genes (Fig. 2). In addition to coding region variations, open chromatin regions in the promoter play a crucial role as a genetic variation resource influencing diversity in grain yield and quality traits among wheat accessions (Fig. 3). These open chromatin regions significantly impact the transcriptional regulation circuit (Fig. 2), particularly by altering TFs binding motifs within these regions. DNA variations in *cis-* motifs likely change the recognition of specific TFs, influencing transcriptional regulation activity (Fig. 3). Indeed, genetic variation within ATAC peaks leads to varied expression of starch and SSP coding genes, ultimately shaping grain yield and quality traits (Fig. 3).

### Comprehensive TRN facilitates systematic identification of regulators governing starch and SSP biosynthesis

Utilizing time-serial RNA-seq and chromatin accessibility data from developing endosperm tissue, we constructed a TRN encompassing starch biosynthesis and SSP accumulation processes (Fig. 4). Among the 436 core TFs within the TRN, we identified numerous functionally characterized factors involved in grain weight or quality, affirming the TRN’s ability to pinpoint crucial regulators for starch and SSP biosynthesis. Notably, a significant portion of the 395 potential novel regulators met individual criteria for functional validation through TILLING mutants, expression-phenotype association analysis in a core collection of wheat accessions, or overlapping analysis with GWAS signals associated with grain size and quality traits. Moreover, 42 TFs appeared in all three screening methods. The collective data reinforces the efficiency and accuracy of our integration strategy in the systematic identification of novel regulators governing starch and SSP biosynthesis in wheat.

### TaABI3-A1, a promising candidate for balanced grain yield and enhanced quality in wheat breeding

In our investigation of 42 high-confidence novel regulators, TaABI3-A1 stood out and underwent detailed functional analysis through generation of knock-out mutants via genome editing. TaABI3-A1 was found to positively activate SSP coding genes, including *TaGlu-1*, *TaLMW-600*, and *TaGli-400*, resulting in lower accumulation of HMW-GS, LMW-GS, and Gliadins in *Taabi3* mutants. Conversely, it inhibits various starch biosynthesis enzyme coding genes, leading to increased TGW and GL when in a loss-of-function state. Notably, in a collection of nearly 300 wheat accessions, an elite haplotype of *TaABI3-A1* was identified, exhibiting higher TGW and grain size. Importantly, this elite haplotype has been selected during the breeding process in China likely with the pursuit of increasing grain yield, especially from 1949 to 1999. The superior of grain yield is associated with alteration of *TaABI3-A1* expression level. Higher TaABI3 expression is linked to lower GL but improved flour quality, as evidenced by increased peak viscosity, breakdown value, and gluten:gliadin ratio. Furthermore, we identified potential upstream regulator TaGBP1 influencing *TaABI3-A1* expression. This suggests the possibility of precisely manipulating the temporal expression level of *TaABI3-A1* to achieve a balance between grain yield and seed quality, especially given the slight temporal differences in starch biosynthesis and SSP accumulation during the grain filling window.

### Valuable data source for mining regulators in shaping grain development

In addition to the *TaABI3-A1* case, we have identified 41 other high-confidence novel candidates with the potential to regulate starch and SSP biosynthesis in wheat. Further research is warranted to thoroughly characterize the functions of these factors. Additionally, exploring natural variations at these gene loci could serve as a valuable resource for breeding improved wheat varieties with superior grain weight and quality, especially when aiming to uncouple the regulation of starch biosynthesis and SSP accumulation. From a broader perspective, the time-course transcriptome and epigenetic dataset as well as the population transcriptome of developing endosperm generated in this study can benefit the research community by facilitating the swift verification of potential candidates. It also provides guidance for studying transcriptional regulatory networks and offers information on development-dynamic open chromatin regions for fine-tuning gene expression, proving useful for precision genome editing in breeding applications.

## Methods

### Plant materials and growth conditions

The wheat cultivar Chinese Spring was used in this study. The seedlings were planted in soil and grown in the greenhouse at 22 °C/20 °C day/night, under long day conditions (16 h light/8 h dark). The stamens were removed before the pollen maturation. Then, we conducted artificial pollination and recorded the number of days to ensure the accurate time of seed development. The seeds at DAP0, DAP2, DAP4, DAP6, DAP8, DAP12, DAP16, and DAP22 stages were sampled for later total RNA extraction and nuclei isolation. The dissection method of endosperm follow the instructions in previous report with some modifications (Zhao et al., 2023). Embryo sacs in DAP0, DAP2 and DAP4 were dissected in a 5% Sucrose solution containing 0.1% RNase inhibitor with fine forceps using the dissecting microscope. For DAP6 and later stages, the relatively independent embryo and seed coat were detached using the same method and dropped, with endosperm left and collected. Endosperm and embryo sacs sampled from about eight spikes were pooled for one biological replicate in early stages and three to five were pooled for one biological replicate in late stages. The RNA-seq (three replicates), ATAC-seq and CUT&Tag (two replicates) experiments at eight or six development stages were carried out.

### Generation and genotyping of *TaABI3 CRISPR/Cas9* lines

To knock out *TaABI3* in wheat *cv.* JM5265, the sgRNA 5’-TCGCCAACTGGATCCTACGG-3’ located in the first exon was used. The CRISPR/Cas9 editing was conducted in Genovo Biotechnology Co. (Beijing, China) following the methods as previously reported (Shan et al., 2014). To identify mutations in *TaABI3-A* (*TraesCS3A02G417300*), *TaABI3-B* (*TraesCS3B02G452200*), and *TaABI3-D* (*TraesCS3D02G412800*), gene-specific primers were designed around the gRNA target site. PCR products were genotyped by Sanger sequencing. Primers used for genotyping are listed in table S12.

### Grain development traits measurement

A Wanshen SC-G seed detector (Hangzhou Wanshen Detection Technology Co., Ltd.) was used to measure grain width, grain length, and TGW, with grains from each plant randomly sampled for three replicates. For starch content measurement, 100 mg flour was used and the content was masured using the Megazyme Total Starch Assay Kit (Megazyme; KTSTA-50A) following manufacturer’s method (three replicates per sample). For the SSP content determination, contents of HMW-GSs, LMW-GSs, and gliadins were detected by RP-HPLC as previously described (three replicates per sample) (Gao et al., 2021). The Agilent Technologies 1260 Infinity IIRP-HPLC system and an Agilent ZORBAX 300SB-C18 column (150 mm × 4.6 mm, 5 μm) were used. Water containing 0.6 mL/L trifluoroacetic acid (solvent A) and acetonitrile containing 0.6 mL/L trifluoroacetic acid (solvent B) were used as elution solvents.

The column temperature was 60 °C and the flow rate was 1 mL/min. The injection volume was 8 μL and the eluent was monitored at 210 nm. The total amounts of HMW-GSs, LMW-GSs, and gliadins were estimated by integrating the relevant RP-HPLC peaks present in the chromatograms.

### Semi-thin sections and coomassie brilliant blue (CBB) staining

Wild-type and *Taabi3* mutant seeds at DAP15 were fixed in FAA solution (63% ethanol, 5% acetic acid, 5% formaldehyde) under vacuum for 2 hours and stayed for 24 hours at room temperature. The samples were dehydrated through a graded ethanol series and embedded in Technovit 7100 resin (Kulzer, https://www.kulzer-technik.de), according to the manufacturer’s instructions. Sections (2-µm thick) were cut using a UC7&2265 microtome (Leica). To visualize starch granules and protein bodies, the sections were stained with 0.1% w/v coomassie brilliant blue R-250 (Tonosaki et al., 2021). Images of stained sections were captured using an OLYMPUS DP74 Microscope. Endosperm starch granules number was manually calculated for six seeds.

### *In situ* hybridization assay

RNA *in situ* hybridization was carried out as described previously (Zhao et al., 2023). Fresh seeds were fixed in formalin-acetic acid-alcohol overnight at 4 °C, dehydrated through a standard ethanol series, embedded in Paraplast Plus tissue-embedding medium (Sigma-Aldrich, P3683), and sectioned at 8 μm width using a microtome (Leica Microsystems, RM2235). Digoxigenin-labeled sense and antisense RNA probes based on the sequence of *TaABI3-A1* were synthesized using a DIG northern Starter Kit (Roche, 11277073910), according to the manufacturer’s instructions. Primers used for the sense and antisense probe synthesis are listed in table S12.

### Luciferase reporter assays

To generate *pTaNAC019-B*: LUC, *pTaARF25*: LUC, *pTaERF3*: LUC, *pTaGlu-1By*: LUC, *pTaGlu-1Dx*: LUC, *pTaLMW-400*: LUC, and *pTaAGPS1a*: LUC, we amplified 2-Kb promoter fragments upstream of each gene from *cv.* Chinese Spring and ligated them with the CP461-LUC as the reporter vector. The ORFs of *TaAG1-A* and *TaABI3-A* were cloned into the Psuper-GFP vector as effectors, and these plasmids were transformed into GV3101 and injected into N. benthamiana leaves in different combinations. Dual luciferase assay reagents (Promega, VPE1910) with the Renilla luciferase gene as an internal control were used for luciferase imaging. The Dual-Luciferase Reporter Assay System kit (Cat#E2940, Promega) was used to quantify fluorescence signals. Relative LUC activity was calculated by the ratio of LUC/REN. The relevant primers are listed in table S12.

### RNA extraction, qRT-PCR analysis, RNA sequencing

Total RNA was extracted using HiPure Plant RNA Mini Kit (Magen, R4111-02). First-strand cDNA was synthesized from 2 μg of DNase I-treated total RNA using the TransScript First Strand cDNA Synthesis SuperMix Kit (TransGen, AT301-02). qRT-qPCR was performed using the ChamQ Universal SYBR qPCR Master Mix (Vazyme, Q711-02) by QuantStudio5 (Applied biosystems). The expression of interested genes was normalized to Actin for calibration, and the relative expression level is calculated via the 2-ΔΔCT analysis method (Livak and Schmittgen, 2001). Primers used for qRT-qPCR are listed in table S12.

Paired-end RNA-seq libraries were prepared and sequenced via the Illumina NovaSeq platform according to the manufacturer’s standard protocols by Annoroad Gene Technology.

### CUT&Tag and ATAC-seq experiment

Embryo sacs and endosperm of DAP0, DAP2, DAP4, DAP6, DAP8, DAP12, DAP16, and DAP22 stages were used to isolate nuclei and carried out the CUT&Tag and/or ATAC-seq experiments. CUT&Tag and ATAC-seq experiments followed the previously described method (Zhao et al., 2023). The fresh samples were used to isolate nuclei, the appropriate number of nuclei (1000-50000) were resuspended by corresponding antibody and secondary antibody. Then the nuclei was incubated with pA-Tn5. *Tn5* transposase used and tagmentation assay is done following the manual (Vazyme, TD501-01). Libraries were purified with AMPure beads (Beckman, A63881) and sequenced using the Illumina Novaseq platform at Annoroad Gene Technology. Antibodies used for histone modifications are listed in table S12.

### Data quality control and alignment

For the analysis of RNA-seq, CUT&Tag, and ATAC-seq data, we first conducted data quality control using the fastp software (Chen et al., 2018). This step ensured the removal of low-quality reads and adapter sequences, resulting in clean datasets for further processing. The cleaned datasets were then aligned to the Chinese Spring reference genome IWGSC RefSeq v1.1 assembly (IWGSC, 2018). For RNA-seq data, the alignment was performed using hisat2 (Kim et al., 2019), while for CUT&Tag and ATAC-seq data, bwa was employed (Li and Durbin, 2009).

### RNA-seq data processing

The raw count of reads of each genes were calculated using FeatureCount software (Liao et al., 2014). And the raw counts were normalized to TPM. The relative expression of each homologous gene in the triad was then normalized to the TPM expression of that gene divided by the total TPM expression of the triad. The identification of balance and bias expression of trads followed previous published strategy (IWGSC, 2018).

Differential gene expression (DEGs) in RNA-seq data was identified using edgeR to compare different samples (Robinson et al., 2010). A threshold absolute value of Log2 Fold Change ≥ 1 and FDR ≤ 0.05 was used for DEGs calling. We employed k-means clustering algorithm for gene clustering, enabling us to categorize genes into meaningful groups based on their expression patterns. Function enrichment analysis were done using an R package ClusterProfiler (Yu et al., 2012).

### Cut&Tag and ATAC-seq data processing

Peak calling for CUT&Tag and ATAC-seq data was conducted using macs2 (Zhang et al., 2008). Given the diverse nature of histone modifications, different parameters were applied for peak calling based on the type of modification. For histone modifications such as H3K27ac and H3K4me3, which are characterized as narrow peaks, we used the parameters “-p 1e-3 -f BAMPE --keep-dup all”. In the case of broad peak histone modifications like H3K27me3, we applied the parameters “—broad --broad-cutoff 0.05 -f BAMPE --keep-dup all”. For ATAC-seq, the peak calling was performed with the parameters “-f BAMPE --keep-dup all --cutoff-analysis --nomodel --shift -100 --extsize 200”. Peaks were assigned to nearest genes using Chipseeker (Yu et al., 2015). The reads under peaks were calculated using FeatureCount (Liao et al., 2014), and the raw counts were normalized to CPM value.

For ATAC-seq data, the TFs binding activity was calculated by R package chromVAR (v1.10.0) (Schep et al., 2017). HINT (Hmm-based IdeNtification of Transcription factor footprints) was used for ATAC-seq footprints identity (Gusmao et al., 2014). JASPAR Plantae database (https://jaspar.genereg.net/) was used as motifs set (Khan et al., 2018). Custome wheat genome were configurated based on the introduction of HINT software using the Chinese Spring reference genome IWGSC RefSeq v1.1.

### Transcriptional regulatory network (TRN) construction and key TFs identification

To construct a robust transcriptional regulatory network, we integrated RNA-seq and ATAC-seq data (Corces et al., 2018; Trevino et al., 2020; Trevino et al., 2021; Zhao et al., 2023). This comprehensive approach allowed us to map transcription factor (TF) interactions and gene regulatory relationships in wheat.

We first used diamond to perform blastp searches, aligning plant transcription factor protein sequences from the JASPAR database with wheat protein sequences which identified TFs-Motif relationships in wheat. For each gene, we analyzed accessible chromatin regions (pACRs) sequences using ATAC-seq footprint analysis which generated the Motif-targets relationships. Combining the TFs-Motif and Motif-targets data, we derived the TFs-targets relationships. Using the GENIE3 algorithm, we analyzed transcriptional correlations between wheat TFs and all genes based on RNA-seq data. Finally, for each transcription factor, we compared the transcriptional correlation of genes with and without the TF footprint in their promoter regions. This comparison was used to calculate the significance of TFs-targets relationships.

We identified core transcription factors (TFs) in wheat endosperm development, focusing on two categories: structure TFs and TFs directly upstream of storage production genes. The TFs significantly enriched in TFs-TFs regulatory network were considered as the structure TFs. The upstream regulators of protein and starch synthesis genes were considered as directly upstream of storage production genes.

### Haplotype analysis of *TaABI3-A1*

Natural variation retrieved from the whole-exome sequencing project of the Chinese wheat mini-core collection (Li et al., 2022), composed of 287 representative selected varieties for the Chinese national collection, were used to assess the allelic variation of *TaABI3-A1*. The polymorphism with missing rate < 0.5, min allele frequency > 0.05, and heterozygosity < 0.5 were retained for further haplotype analysis using Haploview 4.2, and the differences of the grain size phenotypes corresponding to different haplotypes were tested. The haplotype frequency in each breeding process of China and among the major Chinese agro-ecological zones were calculated according to the material information provided (Li et al., 2022).

### Field growth condition and grain development traits analysis of 102 wheat accessions

A total of 102 accessions representing the genetic structure and geographical distribution of a previously characterized common wheat population were selected as a core wheat collection (Wang et al., 2020a). The collection were sown with two replicates, with five 150-cm-long rows for each accession, in field at Beijing (39°54′N, 116°25′E) during crop season 2017-2018. Field experiments were performed using a completely randomized design, and agronomic management followed local practices.

Mature grains of each accession were harvest and air-dried to water content between 11-13%. After moistening for 10 hours, grains were tempered and milled with a Brabender Junior mill (MLU 220, Uzvil, Switzerland) using method AACC 26-21A, and the reserved flour after screened with a 70GG sieve were used to conduct further phenotypic assay of content of starch, gluten, gliadin, and total storage protein, SDS sedimentation volume, pasting characteristics of starch and dough quality parameters.

### Population RNA-seq and transcriptome analysis of 102 wheat accessions

The flowering spikes of each accession were marked with a sign, and six spikes were harvest at DAP10 and DAP20, respectively. The grains in the middle of the spikes were sampled, immediately frozen in liquid nitrogen and then stored at −80°C. The grains of each accession at DAP10 or DAP20 were mixed and divided into two independent biological replicates. Total RNA was extracted using a TRIzol kit (Invitrogen, Carlsbad, CA, USA), and high quality RNA samples detected by the Agilent2100 Bioanalyzer instrument were subjected to PE150 sequencing using a BGISEQ500 platform (BGI, Shenzhen, China). Filtered reads were mapped to the wheat reference genome IWGSC Refseq v1.1 (IWGSC, 2018), and transcripts aligned to each gene were calculated and normalized to TPM values.

### Coefficient of variation and correlation analysis

The SNP frequency and coefficient of variation (CV) was used as a measure of genotypic variability, transcriptional variability and phenotypic variability. Genotypic variability was characterized by SNP frequency. It is calculate using the SNP number divided by the length genomic region, with a percentage of 1% indicates that there is a variation within population every 100 bp. The CV was used to assess the variability of expression level and phenotypic traits, defined as the standard deviation divided by the mean. CV, a unitless statistical measurement independent of size, has been widely used to measure and compare variation of quantitative traits, evolvability and phenotypic plasticity (Pélabon et al., 2020).

To calculate the correlation between gene expression levels and epigenetic modification signals, Pearson correlation coefficients were calculated between the z-scale TPM values and the z-scale CPM values of promoter region (from −3000bp to +1000bp) annotation peaks across each gene. To calculate the correlation between gene expression levels and phenotypic traits, Pearson correlation coefficients were calculated between the gene expression TPM values and trait value of wheat varieties. For the expression correlation between two genes, TPM values in each wheat variety were used to calculate the Pearson correlation coefficients.

### Statistics and data visualization

If not specified, R (https://cran.r-project.org/; version 4.0.2) was used to compute statistics and generate plots. For two groups’ comparison of data, the student’s t-test was used, such as Fig. 3D, 3G, 5G, 6D, 6F, 6G, 6H, 6I, 6J, 7B, and fig. S3C, S3D. For three or more independent groups comparison of data, Fisher’s Least Significant Difference (LSD) was used, such as Fig. 2J, 3A, 3I, 3K, 3L and fig. S3A, S3F. Pearson correlation was used in Fig. 1G, 2C, 2J, 3E, 3H, 3J, 7C, 7E and fig. S2C, S3E.

## Code and Data availability

The raw sequence data of RNA-seq, CUT&Tag and ATAC-seq generated during endosperm development in this study was deposited in the Genome Sequence Archive (https://bigd.big.ac.cn/gsa) under accession number PRJCA022666. The embryo sac data was download from previous study under accession number PRJCA008382 (Zhao et al., 2023).

The data analysis method and code were based on previous study at github (https://github.com/LongZhao1992/Dynamic-chromatin-regulatory-programs-during-embryogenesis-of-hexaploid-wheat.git) (Zhao et al., 2023), and specific data or code are available upon request.

## Author contributions

J.X. designed and supervised the research, J.X., D.-Z. W., L. Z., Z.-H.Z., J.-C.C. wrote the manuscript; L.Z., Z.-H.Z. and D.-Z.W. performed bio-informatics analysis; L.Z., L.-X.L. did ATAC-seq and Cut&Tag experiments; W.-Y. W, D.-Z. W., D.-C.L.did phenotyping and RNA-seq for 102 wheat accessions; D.-Z. W., P. Z. did the haploid and selection analysis; J.-C.C., M.-X G. did *Taabi3* mutants phenotype and transcription regulation assay; X.-L. L. did in situ hybridization; M.-X G., D.-C.L. provides *Taabi3* knock-out transgenic wheats; X.-L. L., D.-C.L., Y.-Y.Y., A.-M.Z. revised the manuscript; D.-Z. W., L. Z., Z.-H.Z., J.-C.C., Y.-M. Y. and J.X. prepared all the figures. All authors discussed the results and commented on the manuscript.

## Acknowledgements

We thank Professor Simon Griffiths (John Innes Centre, UK), Professor Cristobal Uauy (John Innes Centre, UK) and Professor Shifeng Cheng (Guangdong Laboratory for Lingnan Modern Agriculture, China) for sharing the Watkins genotypes and phenotypical data from preprint. We thank Professor Zhensheng Kang (Northwest Agricultural&Forest University, China) and Professor Long Mao (Institute of Crop Science, Chinese Agricultural Academy of Science, China) for sharing some GWAS data and suggestions.

## Supplemental figures and legends

**Fig. S1.**
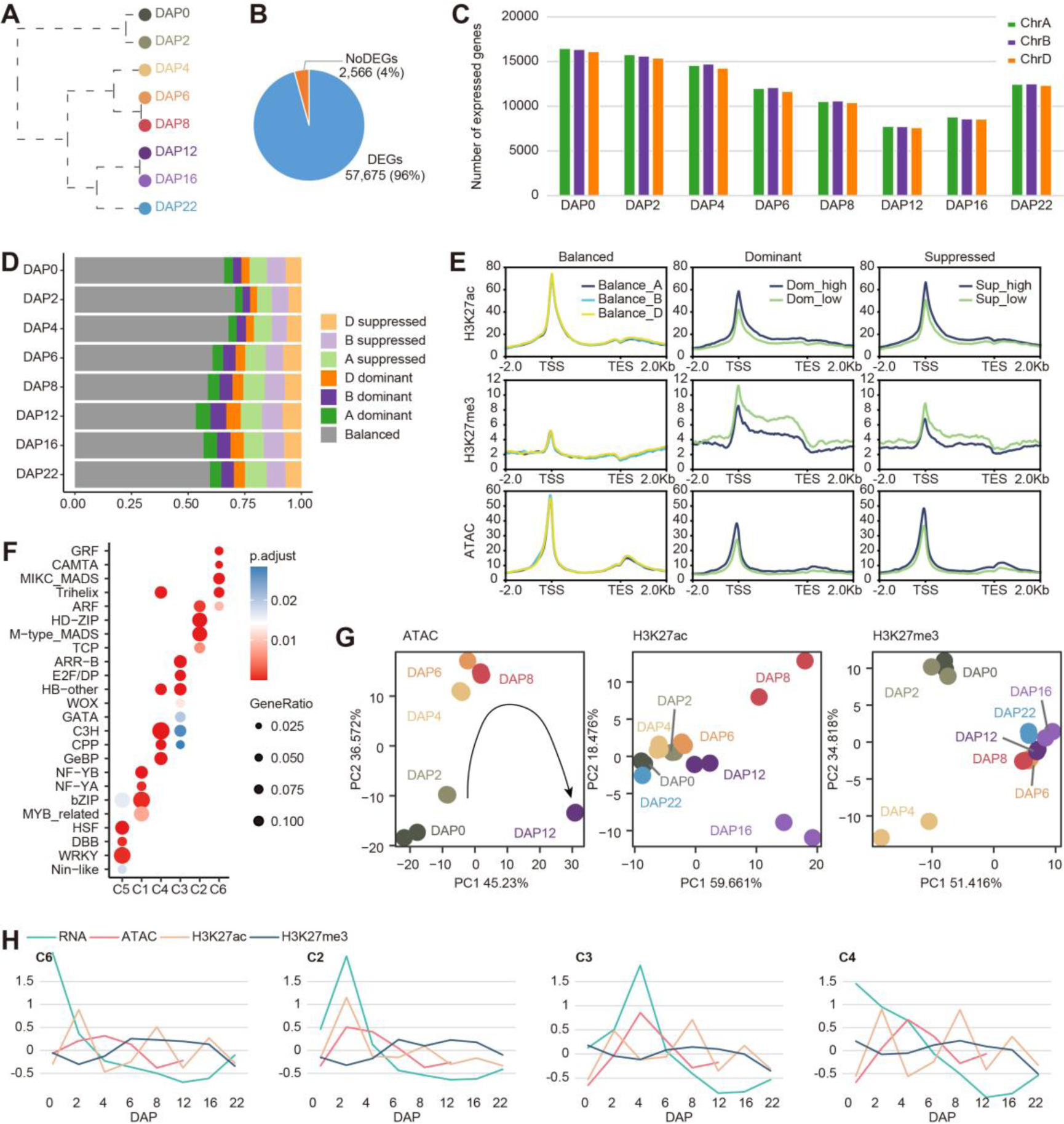
Overview of transcriptome and epigenetic modification during endosperm development in wheat. **A.** Cluster dendrogram of RNA-seq data for DAP0-22. **B.** Proportion of DEGs to total expressed genes in endosperm. The DEGs was calculated between every two samples. **C.** The number of expressed genes in A, B, D sub-genomes at different stages. **D.** Proportion of different subgenome bias expression cluster genes at each sampling stage. **E.** Mega profile of pigenetics modification on bias expression genes. “Dominant high” and “Suppressed high” represent the high expressed allele genes of one traid, while “Dominant low” and Suppressed low” represents the low expressed allele genes of one traid. **F.** TFs families enrichment from different cluster genes. **G.** PCA analysis of ATAC, H3K27ac and H3K27me3. **H.** Correlation between genes’ expression and epigenetics modification in different clusters. Line plot showing the dynamic of Z-score normalized average value of gene expression TPM and epigenetic modification peaks CPM at different DAP.

**Fig. S2.**
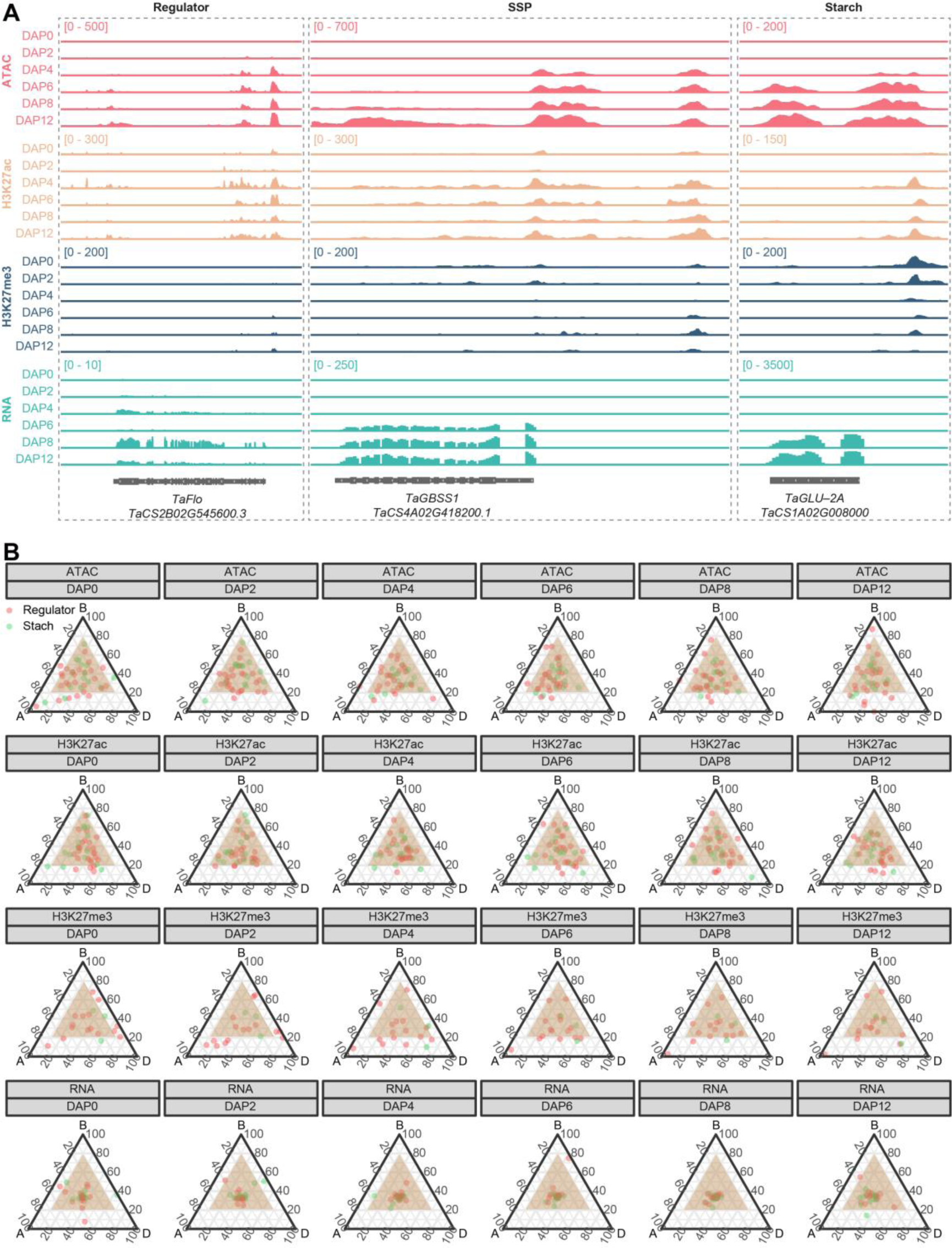
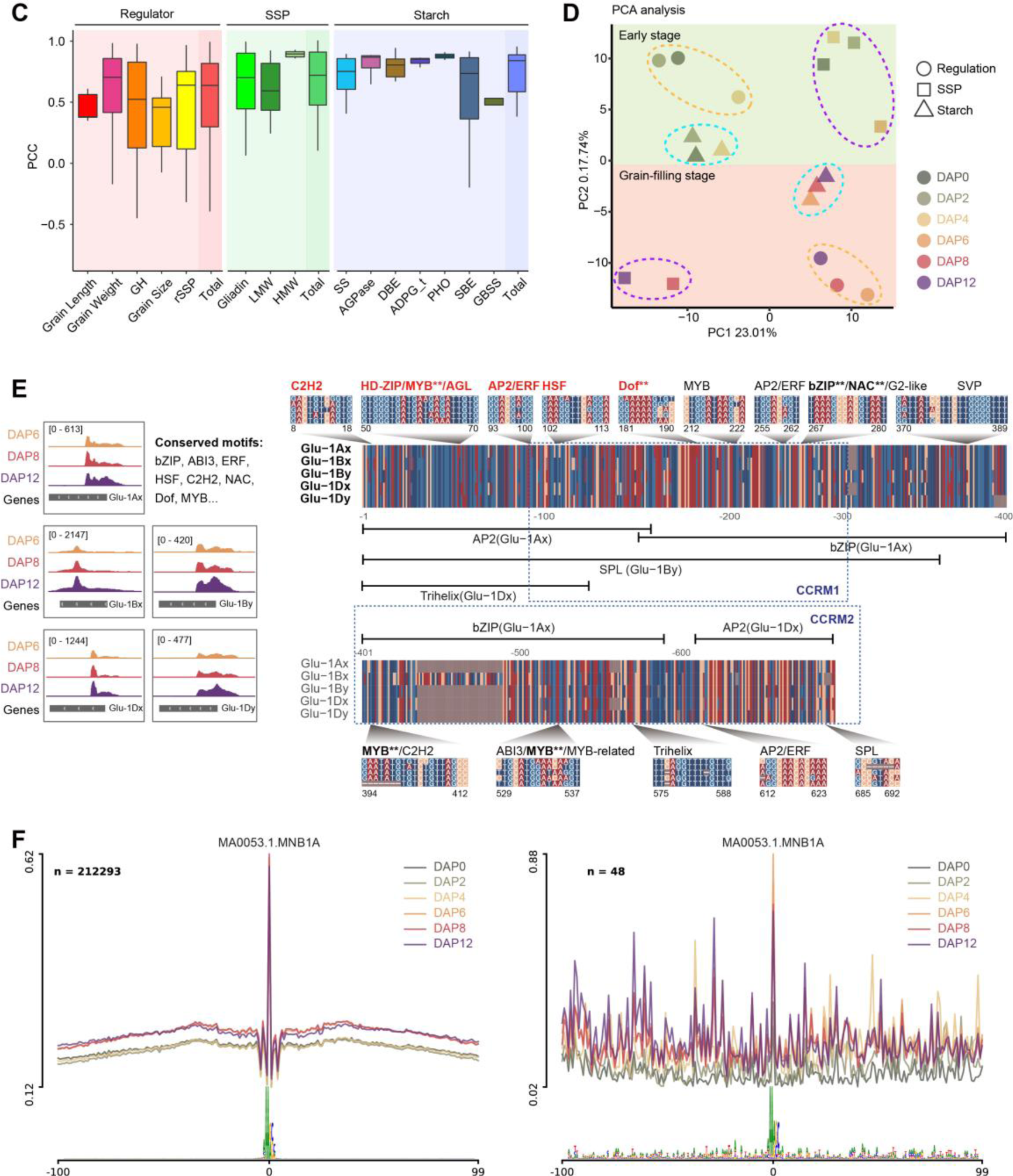
Association of epigenetic modification with starch and SSP coding genes. **A.** Chromatin accessibility, H3K27ac and H3K27me3 profile at representative genes from Starch-, SSP-, and Regulator-genes. **B.** Relative expression and epigenetics modification (ATAC, H3K27ac and H3K27me3) abundance of starch synthesis and regulators genes during endosperm development. Each circle represents a gene triad with an A, B, and D coordinate consisting of the relative contribution of each homoeolog to the overall triad. Balanced triads are shown within brown shadow. **C.** The boxplot of pearson correlation coefficient (PCC) between genes expression across all subgroups and chromatin accessibility at proximal accessible regions (pACRs). PCC values are shown separately for each subgroup and the entire group. **D.** The PCA analysis of variation in TF binding motif activity based on chromVAR. **E.** IGV screenshot showing the similar patterns of ATAC-seq signal around *HMW-GS*, and TF binding motifs in the promoters of *HMW-GS* genes. **F.** The footprint of NMNB1A binding sites in whole-genome level (left) or in SSP coding genes (right) during endosperm development.

**Fig. S3.**
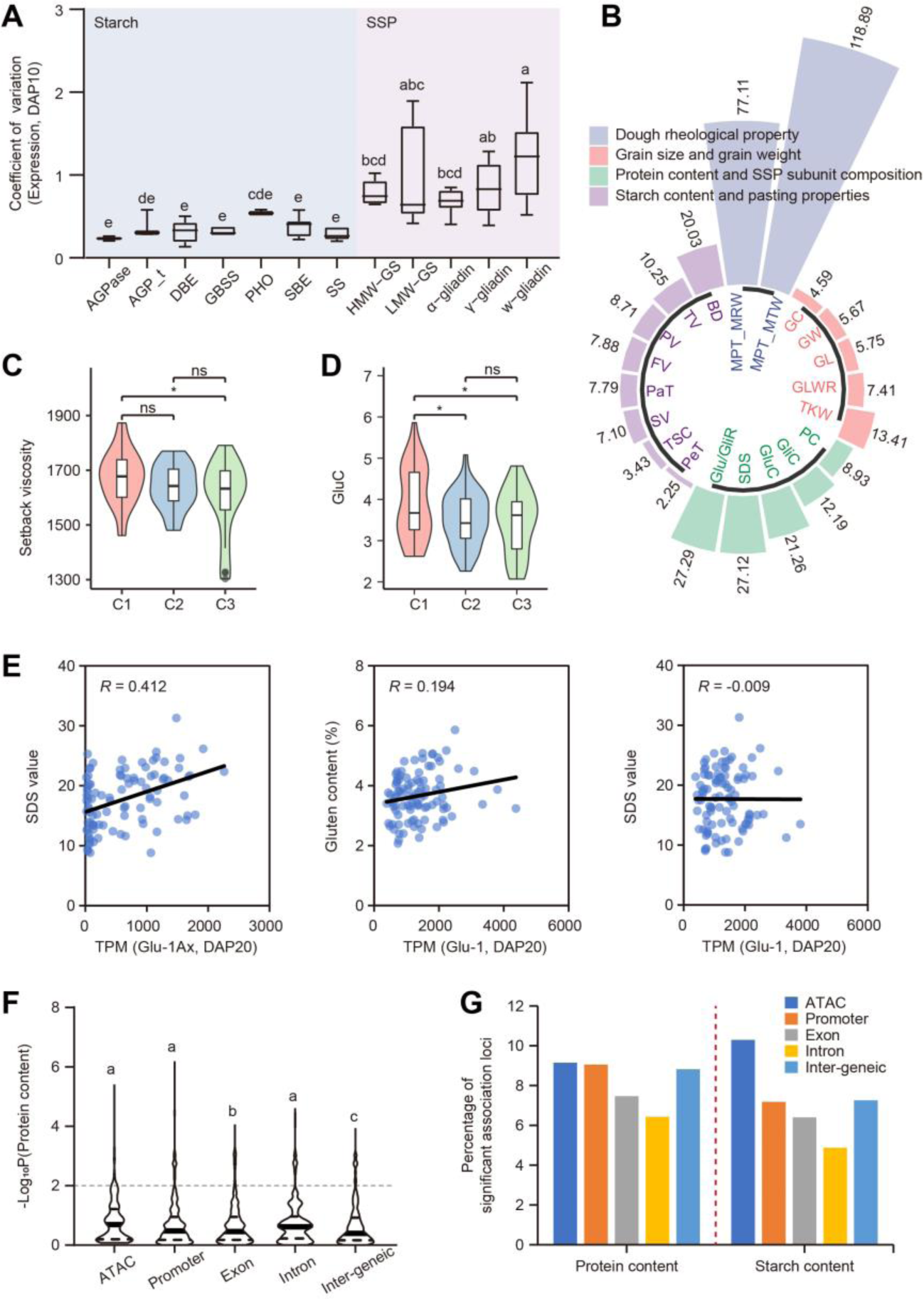
Expression diversity of starch and SSP-related genes within wheat population. **A.** The comparison of coefficient of variation (CV) in expression levels for each subtype genes within the core collection of 102 wheat accession at DAP10. The box denote the 25^th^, median and 75th percentiles, and whiskers indicate 1.5× interquartile range. The LSD multiple comparisons test was used to determine the significance of CV differences among subtype genes. Different letters indicate a significant difference at *P* ≤ 0.05. **B.** Rose plots indicating the CV of each trait across 102 wheat accessions. **C.** Compareison of the setback viscosity (SV) of wheat varieties among three groups clustered by *ADPG_t* expression level. The student’s t-test was used to determine the statistical significance between two groups. *, *p* ≤ 0.05; ns, no significant difference. **D.** Comparison of the gluten contents of wheat varieties among three groups clustered by *Glu-1Ax* expression level. The student’s t-test was used to determine the statistical significance between two groups. *, *p* ≤ 0.05; ns, no significant difference. **E.** Scatter plot of gluten content (left panel) and SDS value (middle panel) against the total expression level of Glu-1, and SDS value against the expression level values of Glu-1Ax (right panel) in corresponding accessions at DAP20. Each dot denotes one accession, and the black line represents the regression trend calculated by the general linear model. **F.** Comparison of the −log_10_(*p* value) of SNPs with storage protein content among different genomic feature. The LSD multiple comparisons test was used to determine the significance differences among subtype genes. Different letters indicate a significant difference at *P* ≤ 0.05. **G.** The percentage of significant association loci with storage protein content and starch content among different genomic feature. The SNP with −log_10_(*p* value) ≥ 2 were considered as significant association loci.

**Fig S4.**
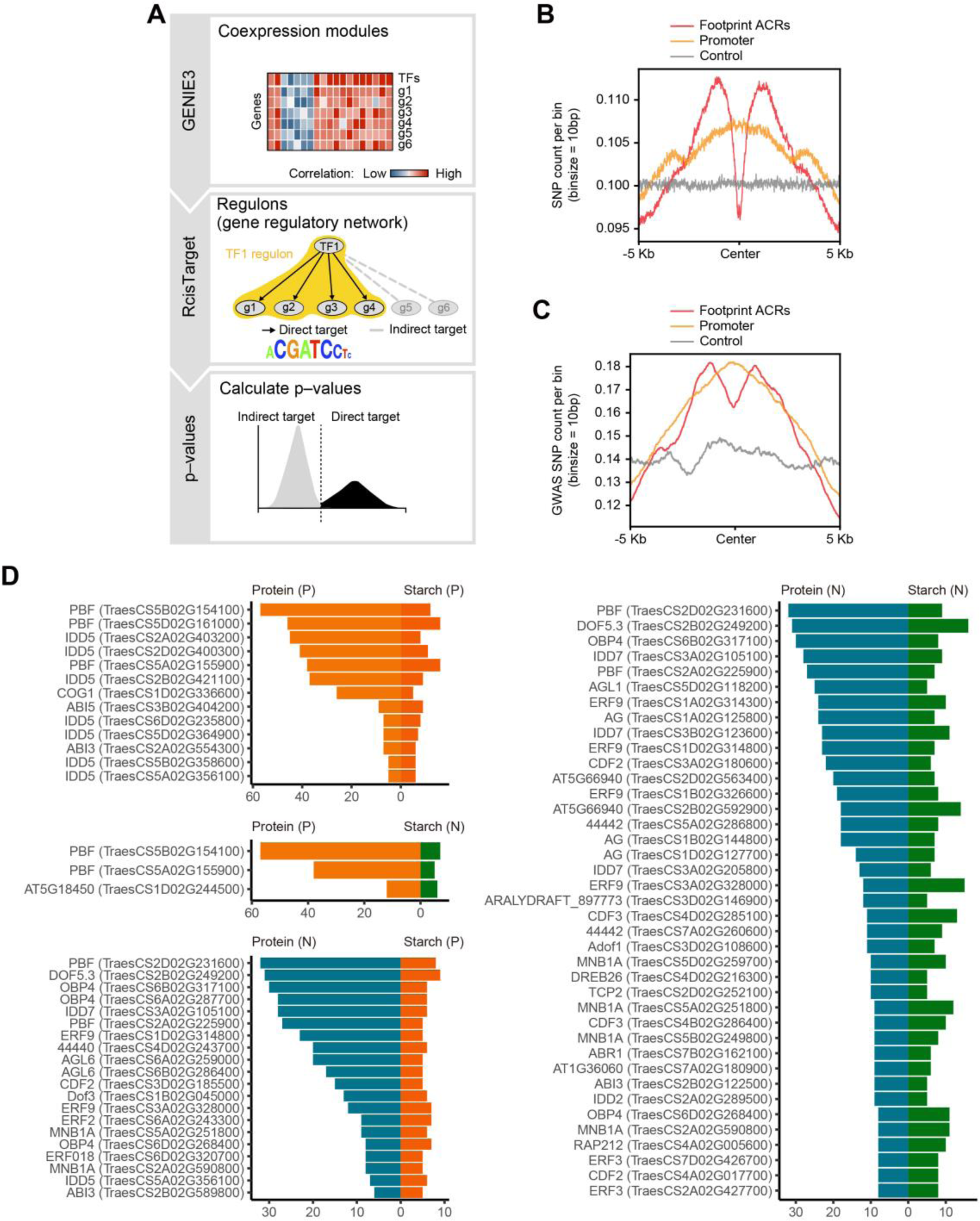
Transcriptional regulatory networks (TRNs) construction. **A**. The flow-chart showing the strategy used for construction of TRNs, according to previous study with minor modification (Aibar et al., 2017). **B, C**. The SNPs (B) and GWAS SNPs (C) distribution around accessible chromatin regions (ACRs) of targets. The footprint ACRs represent the open chromatin regions at the promoter of target genes and can be bound by the TFs. Control represented the random select regions from whole genome. **D**. The number of positively and negatively regulated starch and SSP-related genes for specific TFs.

**Fig. S5.**
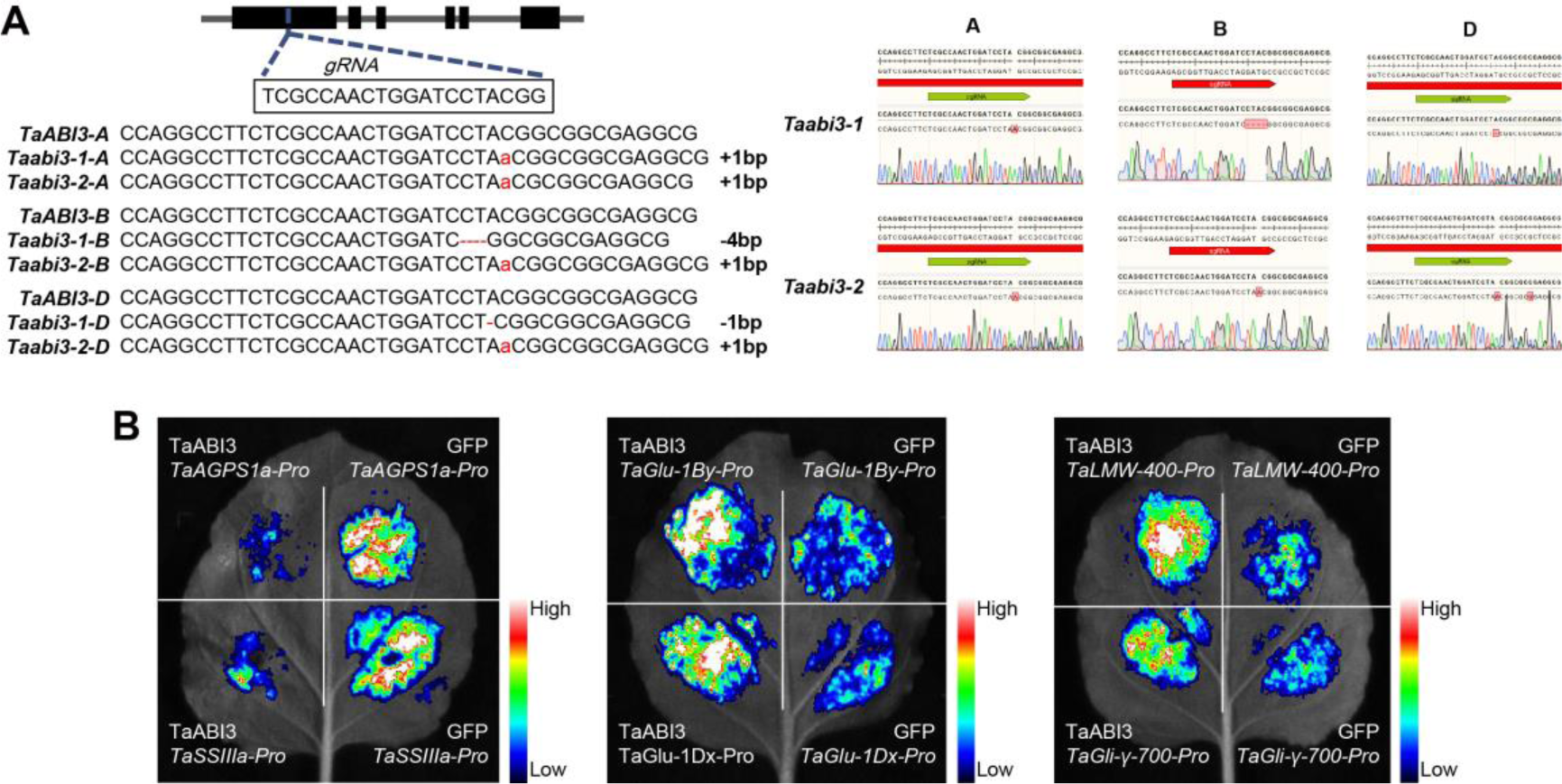
Generation of *Taabi3* mutant lines. **A**. Schematic diagram showing the construction of *Taabi3-KO* lines. The genetic identification of *Taabi3-KO* lines were carried out using Sanger sequencing (right panel). **B**. Representative images for the LUC reporter assay used to evaluate the transcriptional regulation of TaABI3-A1 to its targets involved in starch and SSP biosynthesis.

## Supplemental tables

Supplementary table 1. The TPM values of expressed genes during endosperm development

Supplementary table 2. Clustering of DEGs among developmental stages

Supplementary table 3. The gene expression of SSP coding genes, starch synthesis genes and regulators during endosperm deveploment

Supplementary table 4. The expression level and coefficient of variation of SSP-set, Starch-set genes in wheat population

Supplementary table 5. Correlation between the expression level of SSP-set, Starch-set genes and phenotypic traits in wheat population

Supplementary table 6. Gene information of 436 core TFs from TRN

Supplementary table 7. Grain size and quality GWAS data used in this study

Supplementary table 8. The evaluation of 395 novel TFs

Supplementary table 9. Correlation between the expression level of 395 novel genes and grain size and quality traits in wheat population

Supplementary table 10. The KN9204 TILLING mutants for 395 novel genes

Supplementary table 11. TaABI3-A1 haplotype of MCC population

Supplementary table 12. Primers used in this study

